# Trans- and cis-acting effects of the lncRNA *Firre* on epigenetic and structural features of the inactive X chromosome

**DOI:** 10.1101/687236

**Authors:** He Fang, Giancarlo Bonora, Jordan P. Lewandowski, Jitendra Thakur, Galina N. Filippova, Steven Henikoff, Jay Shendure, Zhijun Duan, John L. Rinn, Xinxian Deng, William S. Noble, Christine M. Disteche

## Abstract

*Firre* encodes a lncRNA involved in nuclear organization in mammals. Here we find that *Firre* RNA is transcribed from the active X chromosome (Xa) and exerts trans-acting effects on the inactive X chromosome (Xi). Allelic deletion of *Firre* on the Xa in a mouse hybrid fibroblast cell line results in a dramatic loss of the histone modification H3K27me3 and of components of the PRC2 complex on the Xi as well as the disruption of the perinucleolar location of the Xi. These features are measurably rescued by ectopic expression of a mouse or human *Firre/FIRRE* cDNA transgene, strongly supporting a conserved trans-acting role of the *Firre* transcript in maintaining the Xi heterochromatin environment. Surprisingly, CTCF occupancy is decreased on the Xi upon loss of *Firre* RNA, but is partially recovered by ectopic transgene expression, suggesting a functional link between *Firre* RNA and CTCF in maintenance of epigenetic features and/or location of the Xi. Loss of *Firre* RNA results in dysregulation of genes implicated in cell division and development, but not in reactivation of genes on the Xi, which retains its bipartite structure despite some changes in chromatin contact distribution. Allelic deletion or inversion of *Firre* on the Xi causes localized redistribution of chromatin contacts, apparently dependent on the orientation of CTCF binding sites clustered at the locus. Thus, the *Firre* locus and its RNA have roles in the maintenance of epigenetic features and structure of the Xi.

## Introduction

X chromosome inactivation (XCI) is controlled by the X-inactivation center that contains the long noncoding RNA (lncRNA) locus *Xist* and some of its regulators such as *Tsix*, *Jpx* and *Ftx* ^1,2^. At the onset of XCI, *Xist* becomes highly expressed on one allele, and the lncRNA coats the future inactive X chromosome (Xi) in cis ^3–5^. Specific proteins are recruited by *Xist* RNA, while others are removed to mediate serial layers of epigenetic modifications, resulting in gene silencing and heterochromatin formation ^6–8^. Epigenetic hallmarks of the Xi include hypo-acetylation of histones H3 and H4, ubiquitination of histone H2A at lysine 119 (H2AK119ubi), de-methylation of histone H3 at lysine 4, tri-methylation of histone H3 at lysine 27 (H3K27me3), di- and tri-methylation of histone H3 at lysine 9 (H3K9me2/3), mono-methylation of histone H4 at lysine 20 (H4K20me), and enrichment in the histone variant macroH2A ^9^. Additional layers of control ensure stability of the silent state of the Xi, including DNA methylation of promoter-containing CpG islands, spatial reorganization of the Xi within the nucleus, and a shift to late replication ^10,11^.

The Xi appears as a heteropycnotic Barr body within the interphase nucleus where it is usually located close to either the nuclear lamina or the periphery of the nucleolus ^12–16^. These two locations are preferred sites of heterochromatin, not only for the Xi but also for other repressed regions of the genome, suggesting that the lamina and the nucleolus play important roles in maintenance of silent chromatin ^12,17^. This is supported by studies in which the perinucleolar space has been proposed to have a primary function in replication and maintenance of repressive chromatin states ^18,19^. The factors and mechanisms that facilitate association of the Xi or of other heterochromatic regions of the genome to specific nuclear compartments such as the lamina or the nucleolus remain elusive. *Xist* RNA interaction with the lamin B receptor (LBR) has been proposed as a critical factor that could help recruit the Xi to the internal rim of the cell nucleus and facilitate chromosome-wide gene silencing ^20^. Our previous studies suggest that perinucleolar positioning the Xi may be facilitated by the lncRNA *Firre* ^21^.

The *Firre* locus comprises conserved tandem repeats essential and sufficient for CTCF binding, which specifically occurs on the Xi but not on the Xa (active X chromosome) ^21–23^. Despite sequence divergence between species, the conserved nature of the repeat locus suggests important roles in mammals. *Firre* RNA is usually confined to the nucleus where it interacts with the nuclear matrix protein hnRNPU ^22,24^. Multiple isoforms of the *Firre* transcript have been reported, including circular RNAs, further complicating an understanding of the roles of *Firre* in different cell types ^25^. On the Xi the *Firre* locus interacts with the *Dxz4* locus, another X-linked repeat that also encodes a lncRNA and binds CTCF only on the Xi ^26–28^. *Dxz4* is necessary for maintenance of the bipartite structure of the Xi ^27,29,30^. Interestingly, the *Firre* locus also interacts with several autosomal regions, consistent with a widespread role in nuclear architecture ^31,32^. A *Firre* knockout (KO) mouse model is viable, but results in cell-specific defects in hematopoiesis that impact common lymphoid progenitors ^31,33^. Importantly, these defects can be rescued by ectopic expression of *Firre* from an autosomal location, thus defining a trans-acting role for *Firre* ^31,33^. RNA-seq analyses of the KO mouse model show organ-specific dysregulation of autosomal gene expression including physiological defects in distinct phases of hematopoiesis ^31^.

Here, we investigate the role of the *Firre* locus and its transcript in maintenance of heterochromatin and 3D structure of the Xi by engineering mouse hybrid cells with allele-specific deletions of the *Firre* locus. Immunostaining studies together with ChIP-seq, CUT&RUN, RNA-seq, ATAC-seq and Hi-C analyses of *Firre* mutants and rescued transgenic cell lines demonstrate trans- and cis-effects of the lncRNA gene.

## Results

### 1. Deletion of Firre on the Xa results in undetectable expression of the lncRNA

Allele-specific CRISPR/Cas9 editing of the *Firre* region was done in Patski cells, in which skewed XCI and frequent species-specific polymorphisms allowed us to design guides to target the Xi from BL6 or the Xa from *Mus spretus* (*spretus*) (Supplementary Table S1). We isolated a single-cell clone with a ~160kb deletion of *Firre* on the Xa (Δ*Firre*^Xa^), a single-cell clone with a ~160kb deletion of *Firre* on the Xi (Δ*Firre*^Xi^) and a single-cell clone with a ~160kb inversion of *Firre* on the Xi (Inv*Firre*^Xi^) (Figure 1A). These genomic alterations and their allele-specificity were confirmed by PCR, Sanger sequencing and DNA-FISH. Deletion of the *Firre* locus on the Xa resulted in undetectable *Firre* expression by RT-PCR (using primers referred to as F/R in Figure 1A), while deletion on the Xi caused no change (Figure 1B, Supplementary Table S2). Allele-specific analysis confirmed the absence of *Firre* RNA-seq reads from either the Xa or the Xi in Δ*Firre*^Xa^ cells, while control loci (*Dxz4*, *Xist*) did not change (Supplementary Table S3).

**Figure 1.**
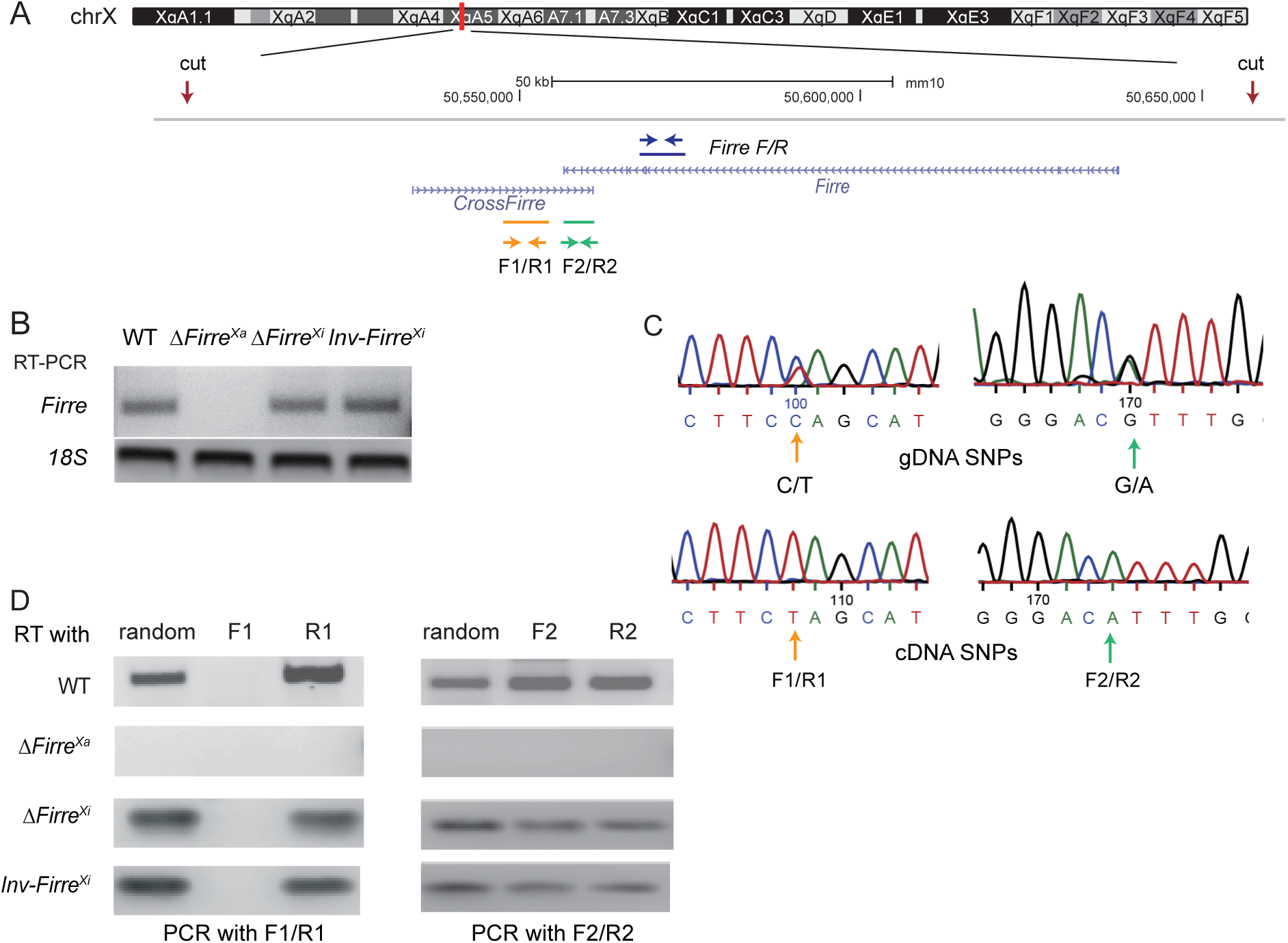
The lncRNAs *Firre* and *CrossFirre* are expressed from the Xa. **A**. Genomic location of *Firre* and *CrossFirre* (GM3561) on the mouse X chromosome (UCSC browser tracks). The location of the CRISPR guide RNAs (cut) used to edit the locus is indicated. The locations of RT-PCR primer pairs to specifically detect *Firre* expression (F/R), strand-specific expression of *CrossFirre* (F1/R1), and strand-specific expression in a region of overlap between *Firre* and *CrossFirre* (F2/R2) are indicated. **B**. RT-PCR analysis using F/R primer pair to detect *Firre* expression in WT, Δ*Firre*^Xa^, Δ*Firre*^Xi^, and Inv*Firre*^Xi^ cells. **C**. Sanger sequencing analysis of *CrossFirre* (F1/R1) and of a region of overlap between *Firre* and *CrossFirre* (F2/R2) confirms heterozygosity of SNPs (BL6 on the Xi and *spretus* on the Xa) in each region assayed. Genomic DNA (gDNA) shows heterozygosity at the SNPs, while the cDNA only shows expression from the *spretus* SNP (Xa). **D**. Strand-specific analysis of *CrossFirre* and *Firre*. Reverse transcription was done using random primers, or F1, R1, F2, R2 primers, followed by PCR using F1/R1 or F2/R2 primer pairs.

*CrossFirre* is an antisense transcript that partially overlaps *Firre* and has been previously reported as imprinted in adult mouse brain (Figure 1A) ^34^. To test its expression pattern in edited Patski cells, we processed strand-specific RT-PCR using a set of primers (F1/R1) that flank a 175bp region in the middle of *CrossFirre* with no overlap with *Firre*, and another set of primers (F2/R2) that flank a 198bp region overlapping the 3’ end of *Firre* (Figure 1A). The presence of SNPs within the two amplified regions was confirmed by Sanger sequencing of genomic DNA (Figure 1C). For the non-overlapped region, we detected forward *CrossFirre* transcripts in WT, Δ*Firre*^Xi^ and Inv*Firre*^Xi^ cells, but these transcripts contained SNPs from *spretus* (Xa) only, and no *CrossFirre* transcripts were detected in Δ*Firre*^Xa^ cells (Figure 1C, D). For the overlapped region, transcripts from both directions were detected in WT, Δ*Firre*^Xi^ and Inv*Firre*^Xi^ cells, but again with SNPs from *spretus* (Xa) only, and with no evidence of either *CrossFirre* or *Firre* transcripts in Δ*Firre*^Xa^ cells (Figure 1D). Using miRNA-seq in either WT or Δ*Firre*^Xa^ we found no evidence of microRNA in the *Firre* region, suggesting that the locus functions independently of the small RNA pathway (Supplementary Table S4).

We conclude that *Firre* and *CrossFirre* are both transcribed from the Xa in Patski cells, since deletion of the entire *Firre* locus on the Xa results in undetectable RNA and its antisense. Note that *Firre* was originally identified as a gene that escapes XCI in both human and mouse ^21,28,32^. However, our current results and those reported in *Firre* KO mouse ES cells and in a *Firre* KO mouse model clearly show that *Firre* is predominantly expressed from the Xa ^31,35^.

### 2. Firre RNA transcribed from the Xa acts in trans to maintain PRC2 and H3K27me3 on the Xi

To determine whether any of the allelic alterations of the *Firre* locus we constructed (Δ*Firre*^Xa^, Δ*Firre*^Xi^, and Inv*Firre*^Xi^) influences epigenetic marks on the Xi, we performed immunostaining analyses of H3K27me3, H2AK119ubi, and macroH2A.1 combined with RNA-FISH for *Xist* to locate the Xi. The majority of nuclei (>95%) had one *Xist* cloud in all cell lines tested, indicating no disruption of *Xist* RNA coating on the Xi (Figure 2A). As expected, a strong H3K27me3 cluster was observed on the Xi in more than 80% of nuclei from WT cells. Surprisingly, Δ*Firre*^Xa^ cells showed a significant reduction by 90% (p <10^-6^, Fisher’s exact test) in the number of nuclei with a H3K27me3 cluster on the Xi (Figure 2B; Table 1). In these nuclei H3K27me3 appeared uniformly mottled throughout the nucleus. In contrast, Δ*Firre*^Xi^ and Inv*Firre*^Xi^ cells showed no significant changes in the number of nuclei with a strong H3K27me3 cluster (80%). No significant changes were observed on the Xi for two other repressive epigenetic modifications known to be associated with XCI, H2AK119ubi and macroH2A.1, when comparing WT and Δ*Firre*^Xa^ (Figure 2C).

**Figure 2.**
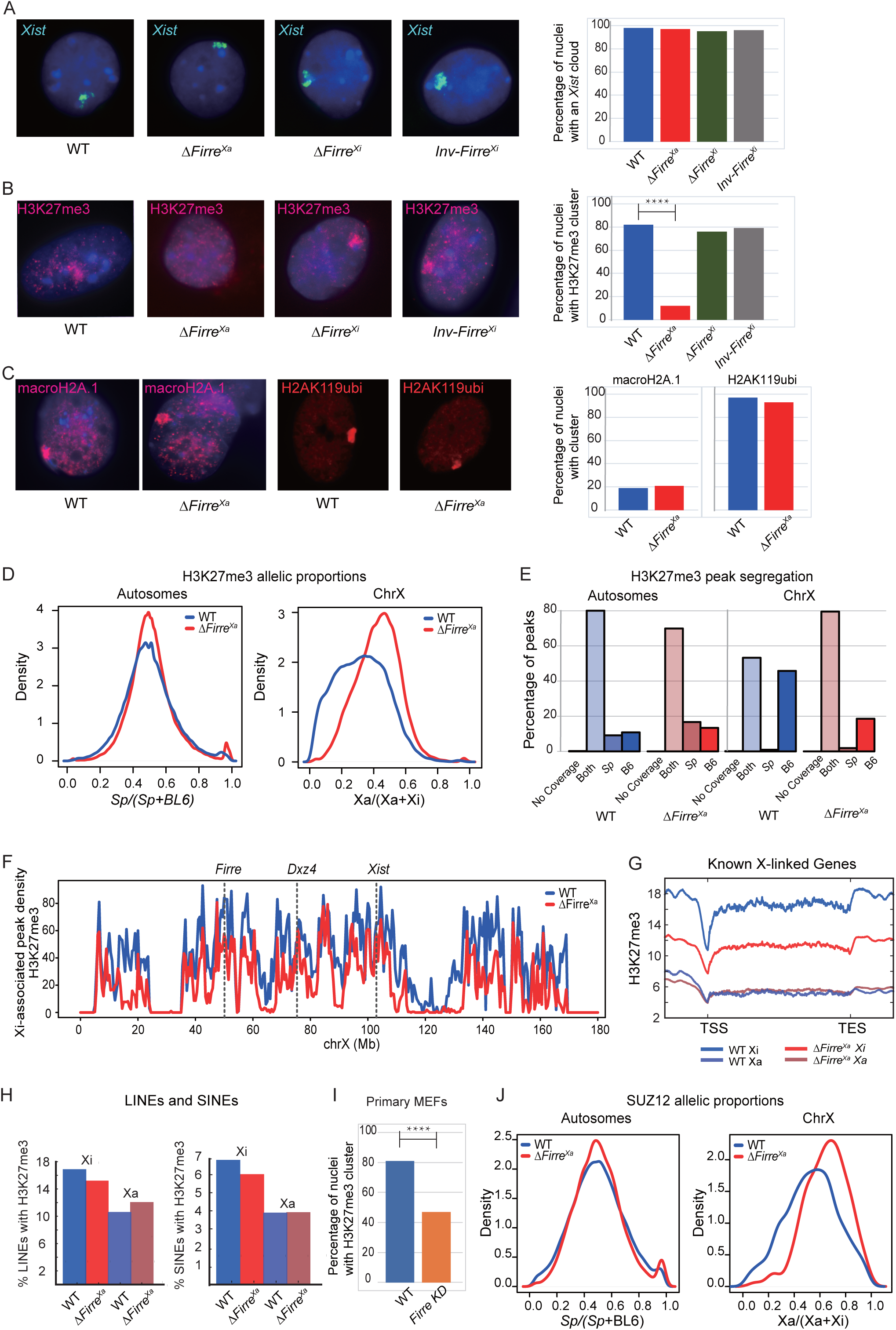
*Firre* RNA transcribed from the Xa acts in trans to maintain H3K27me3 on the Xi in Patski cells and MEFs. **A**. Examples of nuclei after RNA-FISH for *Xist* (green) counterstained with Hoechst 33342 (blue) in WT, Δ*Firre*^Xa^, Δ*Firre*^Xi^ and Inv*Firre*^Xi^ Patski cells, and bar plot of the percentage of nuclei with an *Xist* RNA cloud. No significant difference was detected in the percentage of *Xist* RNA clouds between cell lines by Fisher’s exact test. **B**. Examples of nuclei after immunostaining with an antibody for H3K27me3 (red) counterstained with Hoechst 33342 (blue) in WT, Δ*Firre*^Xa^, Δ*Firre*^Xi^ and Inv*Firre*^Xi^ Patski cells, and bar plot of the percentage of nuclei with a strong H3K27me3 cluster. The percentage of H3K27me3 clusters is significantly lower in Δ*Firre*^Xa^ compared to WT (p <0.00001), while neither Δ*Firre*^Xi^ nor Inv*Firre*^Xi^ significantly differ from WT by Fisher’s exact test. **C**. Examples of nuclei after immunostaining with antibodies for macroH2A.1 (red) and for H2AK119ubi (red) counterstained with Hoechst 33342 (blue), for WT and Δ*Firre*^Xa^, and bar plots of the percentage of nuclei with a strong macroH2A.1 or H2AK119ubi cluster. No significant differences were detected between cell lines by Fisher’s exact test. **D**. Density histograms of the distribution of allelic proportions (*spretus*/(*spretus* + BL6)) of H3K27me3 peaks along autosomes and the X chromosomes for WT (blue) and Δ*Firre*^Xa^ (red). A shift in the distribution of allelic proportions for the X chromosomes due to a decrease in H3K27me3 on the Xi is evident in Δ*Firre*^Xa^ compared to WT (Wilcoxon test: −log10P = inf). **E**. Bar plots of percentages of H3K27me3 peaks in WT (blue) and Δ*Firre*^Xa^ (red) along autosomes and X chromosomes classified as *spretus*-specific, BL6-specific, or at both alleles. **F**. Plots of Xi-associated (common + Xi-specific) H3K27me3 peak density (counts binned within 500kb windows) along the mouse Xi for WT (blue) and Δ*Firre*^Xa^ (red). To account for differences in the number of SNP-covered peaks due to differences in the depth of sequencing between samples, the binned counts are scaled by a factor obtained from the between-sample ratios of autosomal diploid SNP-covered peaks. The position of *Firre*, *Dxz4* and *Xist* is indicated. **G**. Metaplots of average H3K27me3 occupancy at known X-linked genes in WT (Xi blue, Xa purple) and in Δ*Firre*^Xa^ (Xi red, Xa pink). Average enrichment is shown from the transcription start site (TSS) to the termination site (TES), with 3kb (not at scale) on either side. **H**. Bar plots of H3K27me3 enrichment at LINE and SINE repeat regions in WT (Xi blue, Xa purple) and Δ*Firre*^Xa^ (Xi red, Xa pink). **I**. Bar plots of the percentage of nuclei with a H3K27me3 cluster in WT and *Firre* knockdown primary MEFs derived from a WT F1 mouse (BL6 x *spretus*). The percentage of H3K27me3 nuclei with a cluster is significantly lower in cells with a *Firre* knockdown compared to WT MEFs (p <0.0001). **J.** Density histograms of the distribution of allelic proportions (*spretus*/(*spretus* + BL6)) of SUZ12 peaks along autosomes and the X chromosomes for WT (blue) and Δ*Firre*^Xa^ (red). A shift in the distribution of allelic proportions for the X chromosomes due to a decrease in H3K27me3 on the Xi is evident in Δ*Firre*^Xa^ compared to WT (Wilcoxon test: −log10P = 20.98).

**Table 1.**
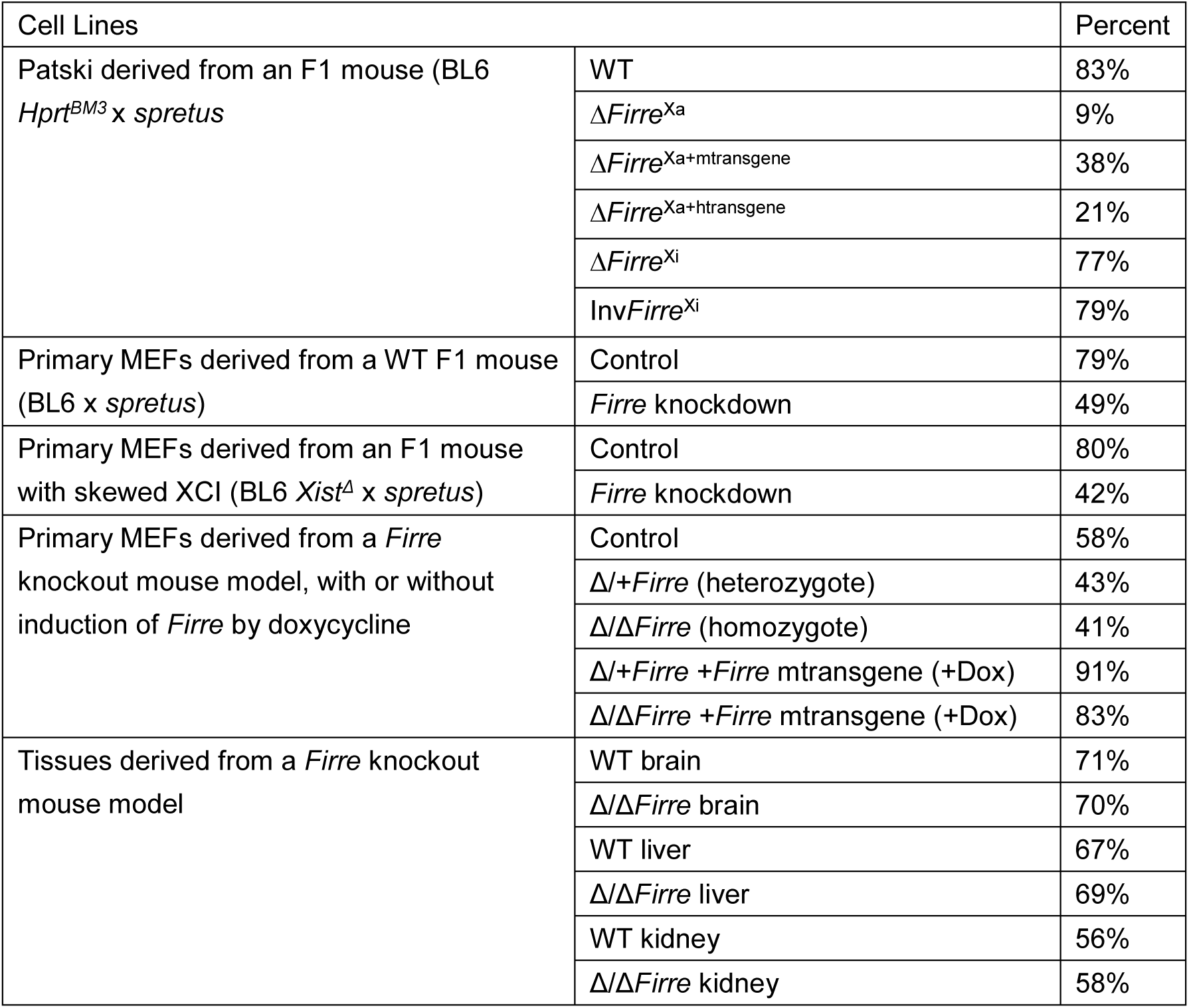
Percentages of cells with a H3K27me3 immunostaining cluster on the Xi

To determine whether H3K27me3 immunostaining intensity decreased specifically on the Xi in Δ*Firre*^Xa^, nuclear regions outside the Xi were examined by ImageJ to quantify fluorescence intensity of these regions using normalization to DNA staining or to histone H4 immunostaining ^36^. This analysis showed no significant difference in H3K27me3 fluorescence intensity between WT and Δ*Firre*^Xa^ cells in regions outside the Xi (Supplementary Figure S1). However, we cannot exclude the possibility of subtle/local changes in the level of H3K27me3 outside the Xi that are undetectable by immunostaining.

The loss of H3K27me3 on the Xi in Δ*Firre*^Xa^ nuclei was confirmed by ChIP-seq, which allowed for a detailed analysis of the distribution of the histone mark throughout the genome. In both WT and Δ*Firre*^Xa^ the distribution of allelic proportions (*spretus*/(*spretus*+BL6)) across H3K27me3 peaks for the autosomes centered close to the anticipated 0.5 reflecting a similar enrichment between alleles (Figure 2D). In contrast, the distribution of allelic proportions for the X chromosomes (Xa/(Xa+Xi)) was centered at ~0.35 in WT cells, consistent with a specific enrichment of H3K27me3 on the BL6 Xi. In Δ*Firre*^Xa^ cells there was a marked shift to higher values (~0.5), consistent with loss of H3K27me3 on the Xi as shown by immunostaining (Figure 2B, D). Diploid H3K27me3 peaks with sufficient SNP read coverage (>5) were designated as either *spretus*-specific or BL6-specific based on an allelic proportion (*spretus*/(*spretus*+BL6)) greater than 0.7 or lower than 0.3, respectively, with the remaining peaks (0.3-0.7) classified as common. Note that the percentage of X-linked H3K27me3 common peaks increased due to a decrease in BL6-specific peaks in Δ*Firre*^Xa^ compared to WT (Figure 2E). Profiles of Xi-associated H3K27me3 peak density demonstrated a chromosome-wide decrease along the Xi in Δ*Firre*^Xa^ compared to WT (Figure 2F). Metagene analyses of X-linked genes confirmed a dramatic decrease of H3K27me3 occupancy along X-linked genes from TSS to TES on the Xi in Δ*Firre*^Xa^ (Figure 2G). Similarly, analyses of LINE and SINE repeats also suggest a partial loss of H3K27me3 in these regions on the Xi in Δ*Firre*^Xa^ (Figure 2H). To extend our observations to a different cell line, we used a double siRNA treatment to knockdown *Firre* in primary WT mouse embryonic fibroblasts (MEFs), which achieved a >67% reduction in level of *Firre* RNA. This resulted in a significant reduction by 43% (p <10^-4^) in the number of nuclei with an intense H3K27me3 staining cluster, supporting our observations in Patski cells (Figure 2I; Table 1).

Next, we investigated SUZ12, a subunit of the PRC2 complex (Polycomb Repressive Complex 2), using CUT&RUN analyses to compare enrichment on the Xi in WT and Δ*Firre*^Xa^. Again, the distribution of allelic proportions (Xa/(Xa+Xi)) for each individual peak showed a pronounced shift toward higher values (~0.75) for the X chromosomes in Δ*Firre*^Xa^ compared to WT, while allelic proportions (*spretus*/(*spretus*+BL6)) for the autosomes were close to the anticipated 0.5 in both samples (Figure 2J). Taken together, these results show that loss of *Firre* RNA leads to a decrease in the PRC2 core component SUZ12, consistent with a decrease in H3K27me3 on the Xi.

We conclude that *Firre* RNA transcribed from the Xa specifically helps target PRC2 for H3K27me3 enrichment on the Xi in trans, but does not influence two other repressive marks associated with XCI, H2AK119ub and macroH2A.1. In contrast, deletion of the *Firre* locus on the Xi has no effect on H3K27me3 staining.

### 3. The loss of H3K27me3 on the Xi can be rescued by a Firre cDNA transgene

To test the possibility of a rescue of the phenotype and confirm a causative role of *Firre* RNA on H3K27me3 enrichment on the Xi, Δ*Firre*^Xa^ cells were transfected with a mouse *Firre* cDNA transgene. As measured by RNA-seq *Firre* expression was restored to a near-normal level after transfection into Δ*Firre*^Xa^ cells. Ectopic expression of the mouse *Firre* cDNA in Δ*Firre*^Xa+mtransgene^ cells rescued the presence of a strong H3K27me3 cluster in 40% of nuclei, supporting a trans-acting role for the RNA (Figure 3A, B; Table 1). Interestingly, ectopic expression of a human *FIRRE* cDNA in Δ*Firre*^Xa+htransgene^ cells also partially rescued the H3K27me3 cluster on the mouse Xi, suggesting partial functional compatibility between species despite sequence divergence (Figure 3A, B; Table 1) ^22^.

**Figure 3.**
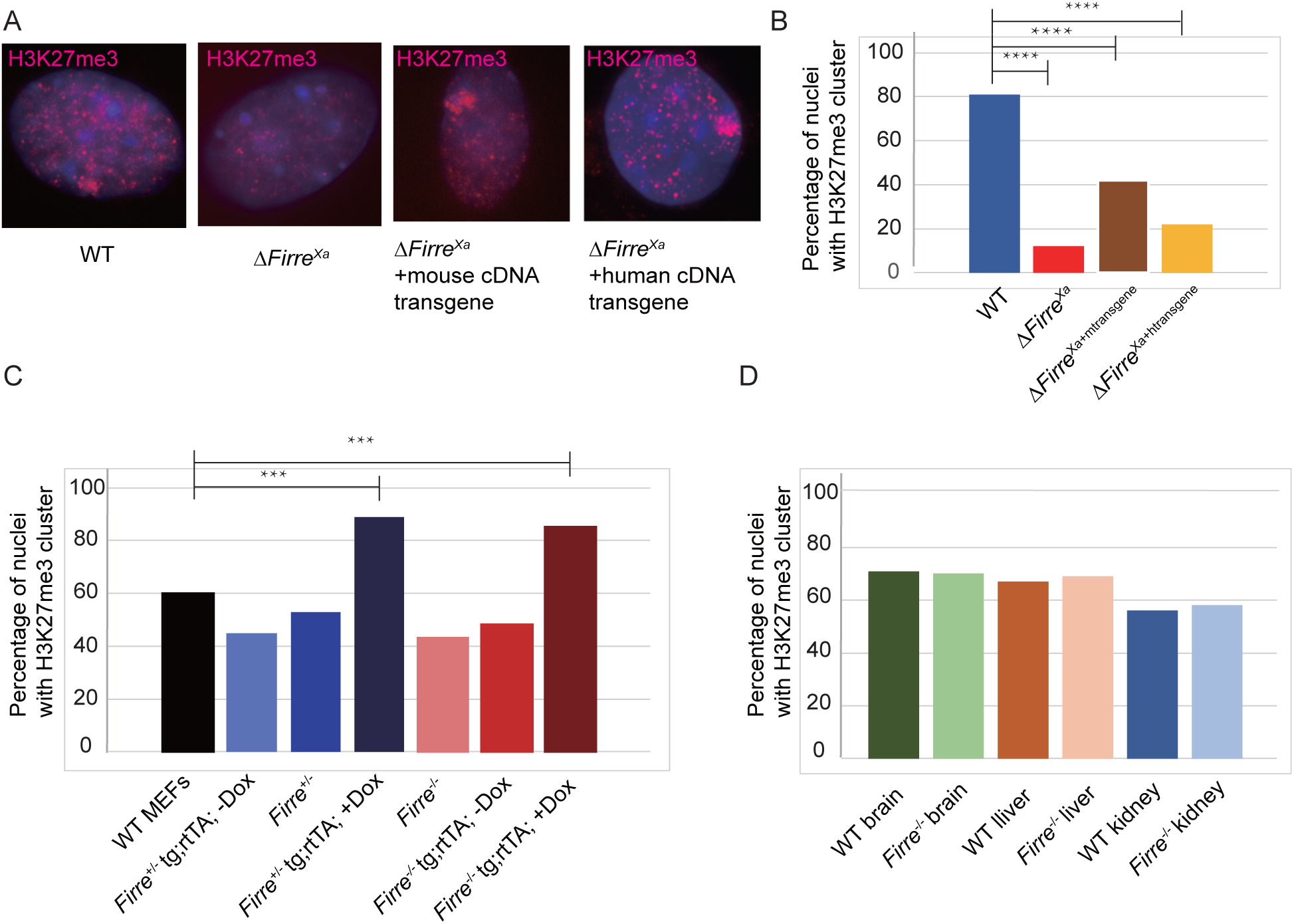
Ectopic expression of *Firre/FIRRE* RNA partly rescues H3K27me3 enrichment on the Xi in Δ*Firre*^Xa^ cells and increases H3K27me3 enrichment in MEFs derived from *Firre* KO mice. **A.** Examples of nuclei after immunostaining with an antibody for H3K27me3 (red) and counterstaining with Hoechst 33342 (blue) in WT, Δ*Firre*^Xa^, and Δ*Firre*^Xa^ transfected with a mouse *Firre* transgene (Δ*Firre*^Xa+mtransgene^) or with a human *FIRRE* transgene (Δ*Firre*^Xa+htransgene^). **B.** Bar plots show the percentage of nuclei with H3K27me3 clusters. The percentage of H3K27me3 clusters in cells transfected with a mouse or human *Firre/FIRRE* transgene significantly increases compared to Δ*Firre*^Xa^ cells by Fisher’s exact test (p <0.0001). **C.** Bar plots of the percentage of nuclei with a H3K27me3 cluster in MEFs derived from WT, heterozygous and homozygous *Firre* KO female mice (*Firre*^+/-^ and *Firre*^+/+^), and in MEFs derived from heterozygous and homozygous KO females that harbor a doxycycline (Dox) inducible transgene (*Firre*+/- tg;rtTA; -Dox; *Firre*+/- tg;rtTA; +Dox; *Firre*-/- tg;rtTA; -Dox; *Firre*-/- tg;rtTA; +Dox). The percentage of nuclei with a H3K27me3 cluster in heterozygous and homozygous *Firre* KO females (*Firre*^+/-^ and *Firre*^-/-^) is lower than in WT MEFs but the difference is not significant (p = 0.0774 for *Firre*^+/-^; p = 0.0567 for *Firre*^-/-^). However, the percentage of nuclei with a H3K27me3 cluster increases significantly in MEFs with an inducible transgene after addition of doxycycline by Fisher’s exact test (p <0.0001). **D.** Bar plots of the percentage of nuclei with a H3K27me3 cluster in tissues derived from WT and *Firre* KO mice show no significant differences (p = 0.9127 for brain, p = 0.8796 for liver, p = 0.762 for kidney).

The role of *Firre* RNA in XCI was further examined in primary mouse embryonic fibroblasts (MEFs) grown from both heterozygous (*Firre* +/- HET) and homozygous (*Firre* -/- KO) female mice from a *Firre* KO mouse model ^33^. H3K27me3 immunostaining of MEFs derived from the *Firre* KO mouse model showed a slight decrease (15-20%) in the proportion of cells with an intense staining cluster on the Xi (Figure 3C; Table 1). Interestingly, induction of *Firre* expression in either heterozygous or homozygous MEFs with an ectopic doxycycline (DOX)-inducible plasmid copy of *Firre* cDNA integrated in the genome resulted in a strong increase in the percentage of cells with a H3K27me3 cluster (>80%), suggesting that ectopically expressed *Firre* RNA can enhance enrichment in H3K27me3 on the Xi in trans in the mouse KO model (Figure 3C; Table 1). H3K27me3 immunostaining in liver, kidney and brain sections derived from homozygous (*Firre* -/- KO) female mice showed no significant decrease of H3K27me3 on the Xi (Figure 3D; Table 1).

Taken together, these observations support a role for *Firre* RNA in maintenance of H3K27me3 on the Xi in differentiated cells. Importantly, a mouse transgene can rescue the loss of H3K27me3 in mutant cells, indicating that RNA from a *Firre* plasmid can perform the trans-effect. Surprisingly, this rescue can also be achieved, at least in part, by a human *FIRRE* transgene, indicating partial conservation of the lncRNA function. The lesser magnitude of changes in H3K27me3 observed in cells from *Firre* KO mice suggests potential compensatory pathways that may act during early development to maintain H3K27me3 on the Xi in KO mice.

### 4. Firre RNA transcribed from the Xa acts in trans to help maintain nuclear location of the Xi

We next examined the effects of allelic *Firre* deletions on the location of the Xi in relation to the nucleolus and the lamina. In WT, Δ*Firre*^Xi^ and Inv*Firre*^Xi^ cells the Xi position was determined by H3K27me3 and nucleophosmin immunostaining to locate the Xi and the nucleoli, respectively (Figure 4A). Since the H3K27me3 cluster is compromised in Δ*Firre*^Xa^, *Xist* RNA-FISH was applied in combination with nucleophosmin immunostaining in Δ*Firre*^Xa^ cells. The Xi location marked by the H3K27me3 cluster or the *Xist* signal was scored as adjacent to either the nucleolus surface, the nuclear periphery, or neither, in a minimum of 200 nuclei. Deletion of *Firre* from the Xa resulted in a significant reduction by 40% (p <0.0001) in the number of nuclei with Xi-nucleolus association and by 50% (p <0.0001) in the number of nuclei with Xi-nuclear periphery association (Figure 4B). Importantly, ectopic expression of a mouse cDNA transgene in Δ*Firre*^Xa+mtransgene^ cells partly rescued Xi-nucleolus and Xi-nuclear periphery associations (Figure 4B; Table 1). Deletion or inversion of *Firre* from the Xi did not alter the location of the Xi (Figure 4A, B; Table 1). Similar analyses in a *Firre* knockdown obtained in MEFs confirmed a decrease in Xi-nucleolus and Xi-nuclear periphery association (Figure 4C; Table 1).

**Figure 4.**
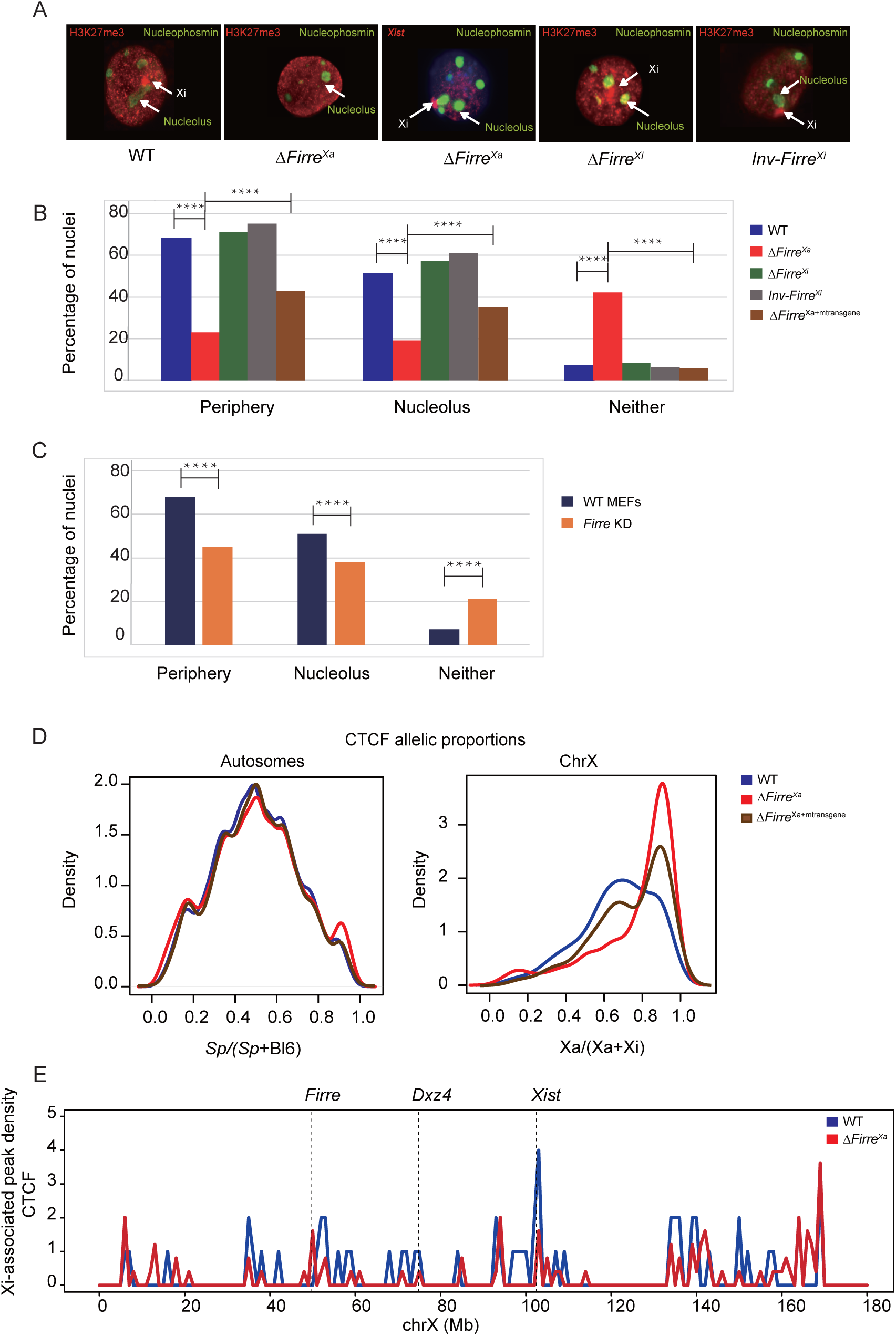
*Firre* RNA transcribed from the Xa acts in trans to maintain Xi location in Patski cells and MEFs. **A.** Examples of nuclei from WT, Δ*Firre*^Xa^, Δ*Firre*^Xi^ and Inv*Firre*^Xi^, after immunostaining for H3K27me3 (red) to locate the Xi in WT, Δ*Firre*^Xi^ and Inv*Firre*^Xi^ or *Xist* RNA-FISH (red) to locate the Xi in Δ*Firre*^Xa^, since there is no H3K27me3 cluster in these nuclei. All nuclei were also immunostained for nucleophosmin (green) to locate the nucleolus. **B.** Bar plots show the percentage of nuclei with the Xi near the periphery, the nucleolus, or at neither in WT, Δ*Firre*^Xa^, Δ*Firre*^Xi^, Inv*Firre*^Xi^ and Δ*Firre*^Xa+mtransgene^ cells. The percentages of periphery- and nucleolus-association of the Xi significantly decrease in Δ*Firre*^Xa^ compared to WT (p <0.0001), but no significant changes are seen in Δ*Firre*^Xi^ and Inv*Firre*^Xi^ by Fisher’s exact test (p = 0.6968). Ectopic expression of the mouse transgene partly rescues Xi location to 41% to the periphery and 34% to the nucleolus. **C.** Bar plots of the percentage of nuclei with the Xi near the periphery, the nucleolus, or at neither in WT and *Firre* knockdown primary MEFs derived from a WT F1 mouse embryo (BL6 x *spretus*). The percentages of periphery- and nucleolus-association of the Xi significantly decrease in *Firre* knockdown compared to control by Fisher’s exact test (p <0.0001). **D.** Density histograms of the distribution of allelic proportions (*spretus*/(*spretus* + BL6)) of CTCF peaks along autosomes and the X chromosomes for WT (blue), Δ*Firre*^Xa^ (red) and Δ*Firre*^Xa+mtransgene^ (brown). A shift in the distribution of allelic proportions due to a decrease in CTCF on the Xi is evident for the X chromosomes in Δ*Firre*^Xa^ compared to WT. In Δ*Firre*^Xa+mtransgene^ the distribution of allelic proportions becomes binomial due to partial restoration of CTCF on the Xi. E. Plots of Xi-associated (common + Xi-specific) CTCF peak density (counts binned within 100kb windows) along the mouse Xi for WT (blue) and Δ*Firre*^Xa^ (red). To account for differences in the number of SNP-covered peaks due to differences in the depth of sequencing between samples, the binned counts are scaled by a factor obtained from the between-sample ratios of autosomal diploid SNP-covered peaks. The position of *Firre*, *Dxz4* and *Xist* is indicated.

CTCF has been implicated in nucleolus association of genomic regions, and we have previously shown that *CTCF* knockdown abolishes Xi-nucleolus association in Patski cells ^21,37^. To further investigate the role of CTCF in positioning the Xi we used CUT&RUN to profile allelic CTCF binding, which showed a strong decrease on the Xi in Δ*Firre*^Xa^ cells (Figure 4D). Indeed, the distribution of allelic proportions (Xa/(Xa+Xi)) for CTCF peaks on the X chromosomes showed a pronounced shift toward higher values (~0.85) in Δ*Firre*^Xa^ compared to WT (~0.65), while allelic proportions (*spretus*/(*spretus*+BL6)) for the autosomes were close to the anticipated 0.5 (Figure 4D). Thus, while there is less CTCF binding on the Xi versus the Xa in WT as expected, CTCF binding is even lower on the Xi in Δ*Firre*^Xa^ (Figure 4E). Interestingly, in Δ*Firre*^Xa+mtransgene^ cells the distribution of allelic proportions for the X chromosomes becomes binomial, suggesting partial restoration of CTCF binding on the Xi (Figure 4D).

We conclude that *Firre* RNA transcribed from the Xa or from an ectopic mouse cDNA transgene can influence in trans the location of the Xi within the nucleus. Loss of *Firre* RNA results in a loss of CTCF binding on the Xi, suggesting potential cooperation between *Firre* RNA and CTCF in maintenance of Xi location.

### 5. Loss of Firre RNA results in changes in gene expression in Patski cells

Next, we examined changes in total gene expression (representing autosomal and X-linked gene expression without discrimination between alleles) in Δ*Firre*^Xa^ cells. About 11% and 14% percent of genes with expression (≥1TPM) in at least one condition were upregulated and downregulated, respectively, in Δ*Firre*^Xa^ compared to WT (Supplementary Table S5). A large proportion of these dysregulated genes were rescued in Δ*Firre*^Xa+mtransgene^ cells (Figure 5A, B, Supplementary Figure S2A, Supplementary Tables S5, S6). Similar results were obtained when considering genes located on autosomes that showed no evidence of copy number changes in WT and Δ*Firre*^Xa^ (Supplementary Figure S2B, Supplementary Table S7). GO analysis showed that the top 20 GO terms for genes upregulated in Δ*Firre*^Xa^ and rescued in Δ*Firre*^Xa+mtransgene^ are related to cell cycle, DNA replication, chromosome segregation, and immune cell function, while a similar analysis for downregulated genes showed they are mainly implicated in development, differentiation and metabolism (Figure 5A, B; Supplementary Table S8). A search for dysregulated genes that may be implicated in histone H3K27 methylation showed that *Ezh1*, *Hist1h1e* and *Hist1h1c* are downregulated genes and *Mki67* is upregulated.

**Figure 5.**
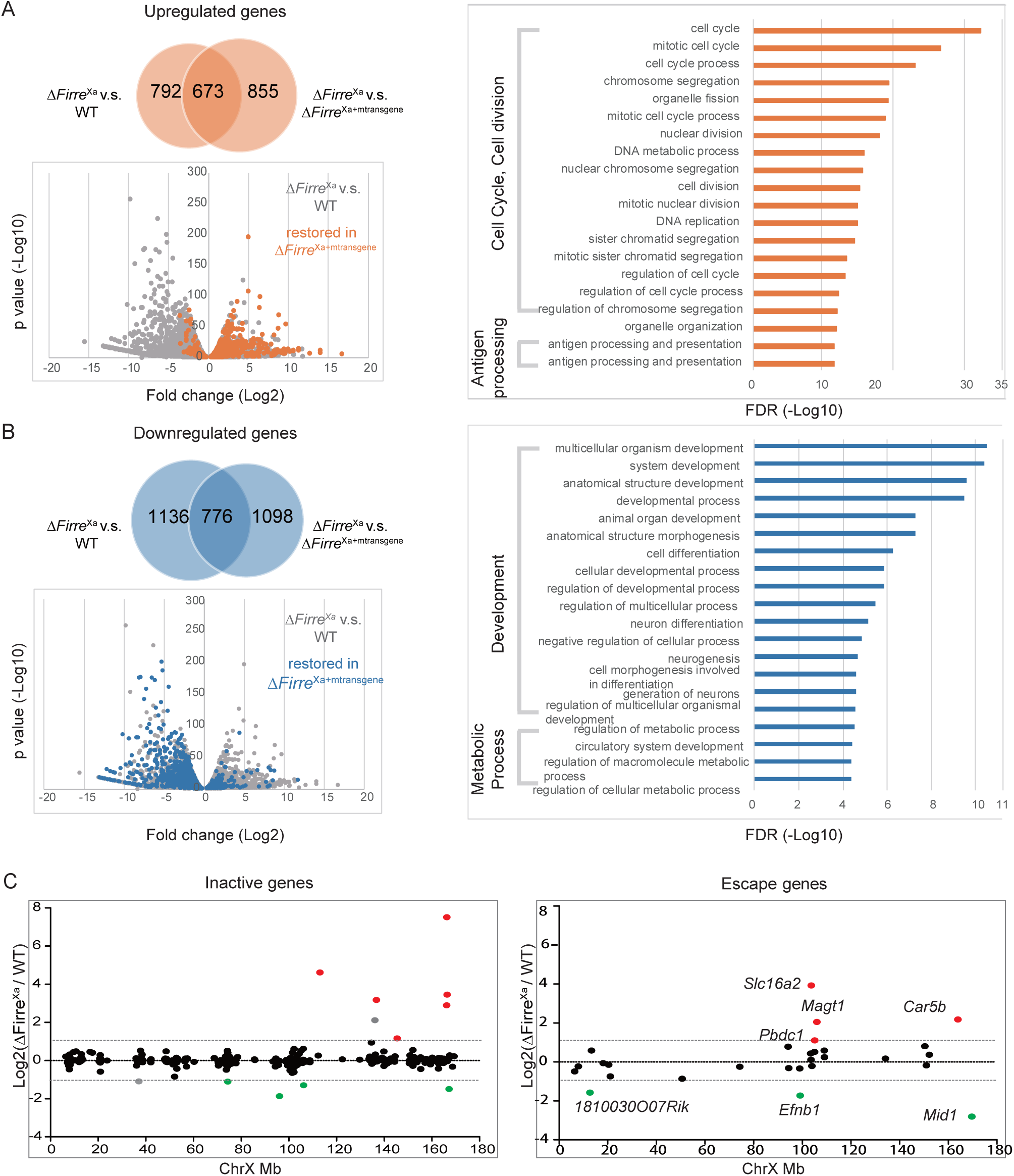
Effects of loss of *Firre* RNA on gene expression. **A**. Upregulated genes in Δ*Firre*^Xa^ cells and Gene Ontology (GO) term enrichment. The Venn diagram shows the number of upregulated genes in Δ*Firre*^Xa^ versus WT and Δ*Firre*^Xa^ versus Δ*Firre*^Xa+mtransgene^. The overlapping gene set represents upregulated genes in Δ*Firre*^Xa^ that are rescued by transgene expression. The scatter plot shows downregulated and upregulated genes in Δ*Firre*^Xa^ versus WT (grey), with genes rescued by reduced expression in Δ*Firre*^Xa+mtransgene^ versus Δ*Firre*^Xa^ (more than 2-fold; p-value < 0.05) highlighted in orange. The top 20 GO terms of overlapping upregulated genes in Δ*Firre*^Xa^ versus WT and rescued in Δ*Firre*^Xa+mtransgene^ are listed. The X-axis indicates the FDR (-log10). **B**. Downregulated genes in Δ*Firre*^Xa^ cells and Gene Ontology (GO) term enrichment. The Venn diagram shows the number of downregulated genes in Δ*Firre*^Xa^ versus WT and Δ*Firre*^Xa^ versus Δ*Firre*^Xa+mtransgene^. The overlapping gene set represents downregulated genes in Δ*Firre*^Xa^ that are rescued by transgene expression. The scatter plot shows downregulated and upregulated genes in Δ*Firre*^Xa^ versus WT (grey), with genes rescued by increased expression in Δ*Firre*^Xa+mtransgene^ versus Δ*Firre*^Xa^ (more than 2-fold; p-value < 0.05) highlighted in blue. The top 20 GO terms of overlapping downregulated genes in Δ*Firre*^Xa^ versus WT and rescued in Δ*Firre*^Xa+mtransgene^ are listed. The X-axis indicates the FDR (-log10). **C**. Xi-expression fold changes for genes that are subject to or escape XCI between Δ*Firre*^Xa^ and WT cells. X-linked genes were grouped based on XCI status. Upregulated genes are colored in red and downregulated genes are colored in green. To note, one upregulated gene and one gene subjected to XCI are colored grey, as they showed more than 2-fold change in expression but with p value > 0.05 in Δ*Firre*^Xa^ versus WT. Genes are ordered (left to right) from centromere to telomere along the Xi.

To determine whether X-linked gene expression was specifically disrupted upon loss of *Firre* RNA we evaluated allelic gene expression. On the Xi only 6/352 genes known to be subject to XCI show >2-fold upregulation in Δ*Firre*^Xa^ cells, suggesting that *Firre* RNA depletion has a minor effect on gene reactivation on the Xi (Figure 5C, Supplementary Tables S3, S5). Four of the reactivated genes are located at the distal end of the Xi where higher accessibility and decreased contact density is observed by ATAC-seq and Hi-C (see below). Although their number is small, a greater proportion of genes that escape XCI (10-14%) were dysregulated compared to genes subject to XCI (2-3%) (Figure 5C, Supplementary Table S5). On the Xa about 4-8% of genes were dysregulated (Supplementary Figure S2C, Supplementary Tables S3, S5). A majority of dysregulated X-linked genes (70-77%), including all 4 upregulated escape genes, were rescued in Δ*Firre*^Xa+mtransgene^ (Supplementary Figure S2D, Supplementary Tables S3, S5).

Next, we investigated whether gene dysregulation may reflect changes in epigenetic features. Metagene plots of genes grouped based on their expression changes in Δ*Firre*^Xa^ and Δ*Firre*^Xa+mtransgene^ together with profiles of individual genes showed a slight increase in H3K27me3 for downregulated genes, and very little or no change in H3K27me3 for upregulated genes or for genes with unchanged expression (Supplementary Figure S2E, F). Note that 90% of dysregulated genes have no H3K27me3 enrichment in either WT or Δ*Firre*^Xa^. Using CUT&RUN analysis for two active epigenetic marks, H3K36me3 and H3K4me3, we found no significant changes on the Xi in Δ*Firre*^Xa^ compared to WT, consistent with little Xi reactivation (Supplementary Figure S3A, B).

Thus, the loss of *Firre* RNA causes changes in gene expression, which are in large part rescued by a transgene. Changes in gene expression may indirectly reflect changes in cell growth and physiology and are not associated with major epigenetic changes. Very little reactivation occurs on the Xi, despite the observed loss of H3K27me3.

### 6. Allelic alterations of Firre cause localized changes in Xi structure

Chromatin accessibility was measured by ATAC-seq. As expected, in WT cells the distribution of autosomal allelic proportions (*spretus*/(*spretus*+BL6)) centered at 0.5, while the X-chromosome allelic proportions (Xa/(Xi+Xa)) were skewed towards the Xa (0.95), indicative of fewer ATAC peaks and lower chromatin accessibility on the Xi. This pattern did not change in Δ*Firre*^Xa^ cells, indicating no overall change in chromatin accessibility on the Xi, consistent with near absence of reactivation (Figure 6A, B). Plots of Xi-associated ATAC peak density in Δ*Firre*^Xa^ captured the low accessibility profile of the Xi in both WT and Δ*Firre*^Xa^. However, a higher peak density was observed in the telomeric region of the Xi in Δ*Firre*^Xa^ where some reactivated genes are located (Figure 5C; 6C). ATAC-seq patterns and allelic proportions were unchanged in Δ*Firre*^Xi^ and Inv*Firre*^Xi^ cells (Supplementary Figure S4A, B). To determine whether *Dxz4* and *Firre* may have synergistic cis-effects on chromatin accessibility on the Xi, ATAC-seq was done on a double-mutant line Δ*Firre*^Xi^/ΔDxz4^Xi^. Interestingly, a pronounced shift to lower values in the distribution of allelic proportions for the X chromosomes (peak at ~0.55) was observed in the double mutant compared to ΔDxz4^Xi^ (peak at ~0.85) and to Δ*Firre*^Xi^ (peak at ~1), indicating increased chromatin accessibility on the Xi in the double mutant (Figure 6D).

**Figure 6.**
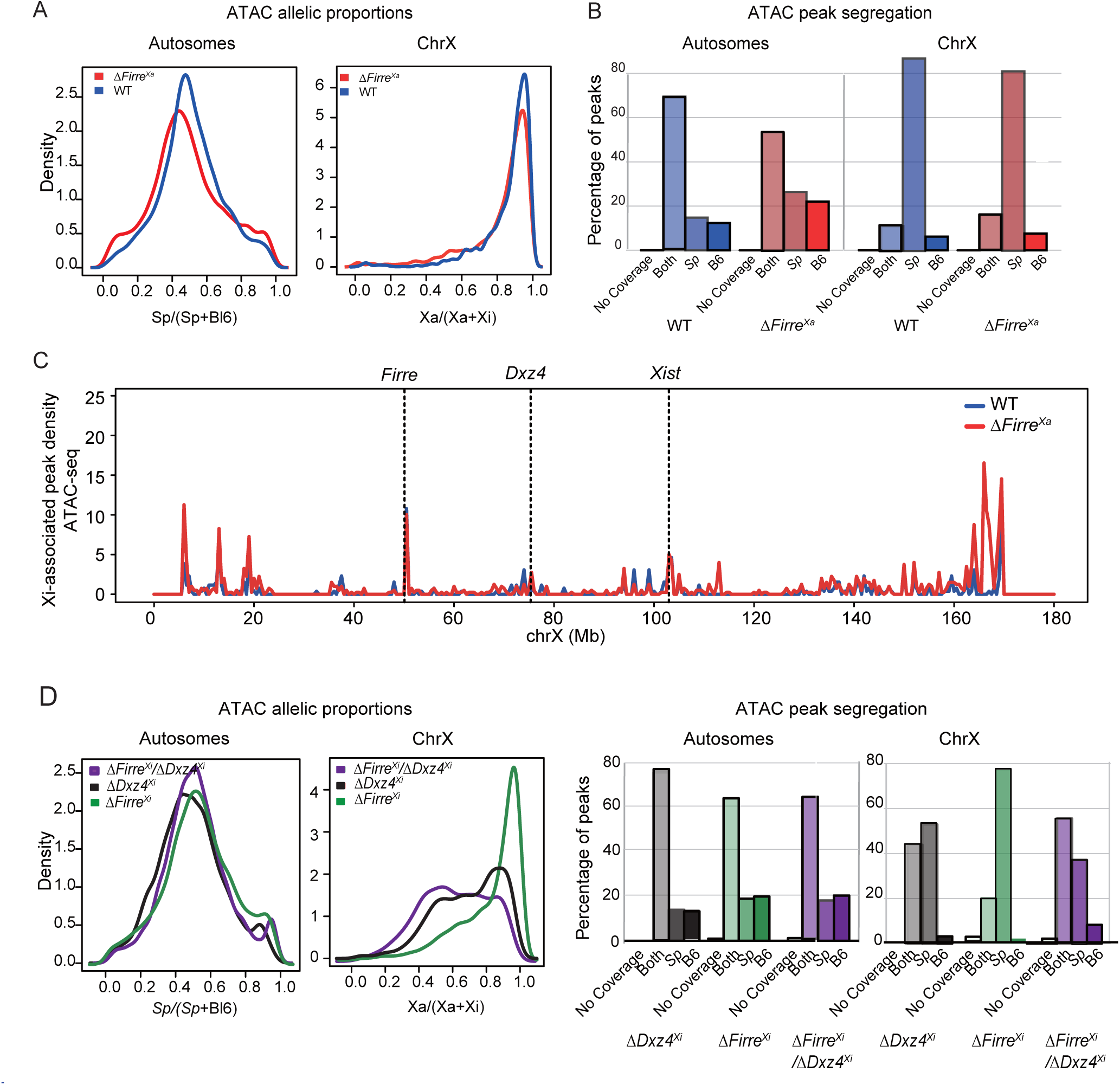
Effects of allelic deletions of *Firre* and of a double *Firre/Dxz4* deletion on chromatin accessibility. **A**. Density histograms of the distribution of allelic proportions (*spretus*/(*spretus* +BL6)) of ATAC peaks along the autosomes and the X chromosomes for WT (blue) and Δ*Firre*^Xa^ (red). No shift is observed (Wilcoxon test: −log10P = 32). **B**. Percentages of ATAC peaks in WT (blue) and Δ*Firre*^Xa^ (red) along the autosomes and the X chromosomes classified as *spretus*-specific, BL6-specific, or at both. **C**. Plots of Xi-associated (common + Xi-specific) ATAC peak density (counts binned within 500 kb windows) along the Xi for WT (blue) and Δ*Firre*^Xa^ (blue). To account for differences in the number of SNP-covered peaks obtained between samples due to differences in the depth of sequencing, the binned counts are scaled by a factor obtained from the between-sample ratios of autosomal diploid SNP-covered peaks. The position of *Firre*, *Dxz4* and *Xist* is indicated. **C**. Density histograms of the distribution of allelic proportions (*spretus*/(*spretus* +BL6)) of ATAC peaks along the autosomes and X chromosomes in Δ*Dxz4*^Xi^ (black), Δ*Firre*^Xi^ (green) and the double mutant Δ*Firre*^Xi^/Δ*Dxz4*^Xi^ (purple). A shift to a lower Xa/(Xa+Xi) ratio is observed in the double mutant compared to Δ*Dxz4*^Xi^ and Δ*Firre*^Xi^, consistent with increased accessibility on the Xi (Wilcoxon test: −log10P = 35). **D**. Percentages of ATAC peaks in Δ*Dxz4*^Xi^ (black), Δ*Firre*^Xi^ (green), and the double mutant Δ*Firre*^Xi^/Δ*Dxz4*^Xi^ (purple) along the autosomes and the X chromosomes classified as *spretus*-specific, BL6-specific, or at both.

In situ DNase Hi-C was done to generate high-resolution allele-specific contact maps in cells with alterations at the *Firre* locus. Δ*Firre*^Xa^ cells retained the characteristic bipartite structure of the Xi, but changes were detected in contact distribution (Figure 7A, Supplementary Figure S5A). Specifically, contacts in the centromeric superdomain (ChrX:5-75Mb) were attenuated, while contacts in a region within the telomeric superdomain between *Dxz4* and *Xist* (ChrX:75-100Mb) were strengthened. Interestingly, a loss of contacts was observed in the distal telomeric region (ChrX:165-170Mb) of the Xi, corresponding to a region that contains reactivated genes with increased chromatin accessibility as described above (Figures 5C, 6C, 7A). A previous study reported that the strong TAD boundary located at *Firre* is preserved but weakened upon deletion of the locus in MEFs ^23^. Our current allelic analyses show a strong boundary at or close to the *Firre* locus on the Xi and a weaker boundary on the Xa in WT cells, the latter being abrogated on the Xa but not on the Xi in Δ*Firre*^Xa^ (Figure 7B, Supplementary Figure S5C). Insulation score analysis confirms loss of insulation around *Firre* only on the Xa in Δ*Firre*^Xa^ (Figure 7C). In contrast, the TAD boundary around the *Firre* locus is maintained upon deletion and inversion of the locus on the Xi (Supplementary Figure S5C).

**Figure 7.**
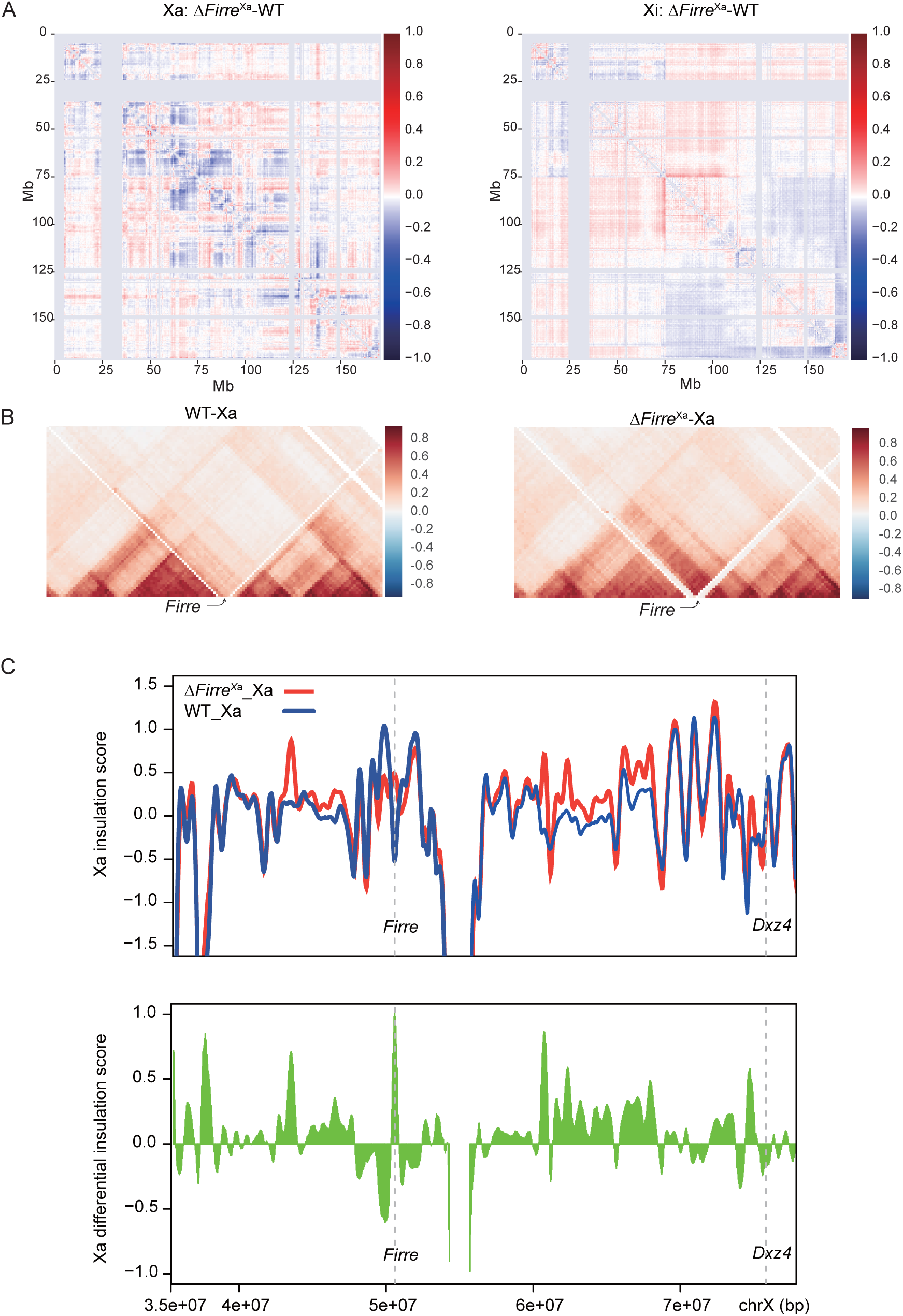
Loss of *Firre* RNA from the Xa causes localized redistribution of contacts on the Xa and Xi. **A.** Pearson correlated-transformed differential contact maps of the Xa and Xi at 500kb resolution to highlight differences between Δ*Firre*^Xa^ and WT. Loss and gain of contacts in the Δ*Firre*^Xa^ versus WT appear blue and red, respectively. The color scale shows differential Pearson correlation values. **B**. Pearson correlated-transformed contact maps (40kb resolution) of the Xa for 4Mb around the *Firre* locus highlight the loss of the strong boundary between TADs on the Xa in Δ*Firre*^Xa^ versus WT. See also Supplementary Figure S4B for corresponding maps of the Xi. **C**. Insulation score profiles at 40kb resolution for the whole Xa in WT (blue) and Δ*Firre*^Xa^ (red). A differential insulation score profile of the Xa (green) in Δ*Firre*^Xa^ relative to WT based on 40kb resolution is shown below. The position of *Firre* and *Dxz4* is indicated.

In Δ*Firre*^Xi^ cells the Xi bipartite structure was intact, but there was an increase in contacts within the centromeric (ChrX:50-75Mb) and telomeric (ChrX:75-100Mb) superdomains, and between the two superdomains, suggesting that the *Firre* locus, though not essential, acts in cis to help shape the bipartite structure of the Xi (Figure 8A, Supplementary Figure S5B). Similarly, Inv*Firre*^Xi^ cells showed persistence of the bipartite structure, but also a redistribution of contacts around the *Firre* locus (Figure 8A, Supplementary Figure S5B). By virtual 4C we observed a loss of contacts between *Firre* and the centromeric region (ChrX:42-50Mb) and a gain of contacts between *Firre* and *Dxz4* (ChrX:50-75 Mb), suggesting that *Firre* contacts along the Xi are orientation-dependent (Figure 8B), reminiscent to the orientation-dependent contacts made by *Dxz4* ^29^.

**Figure 8.**
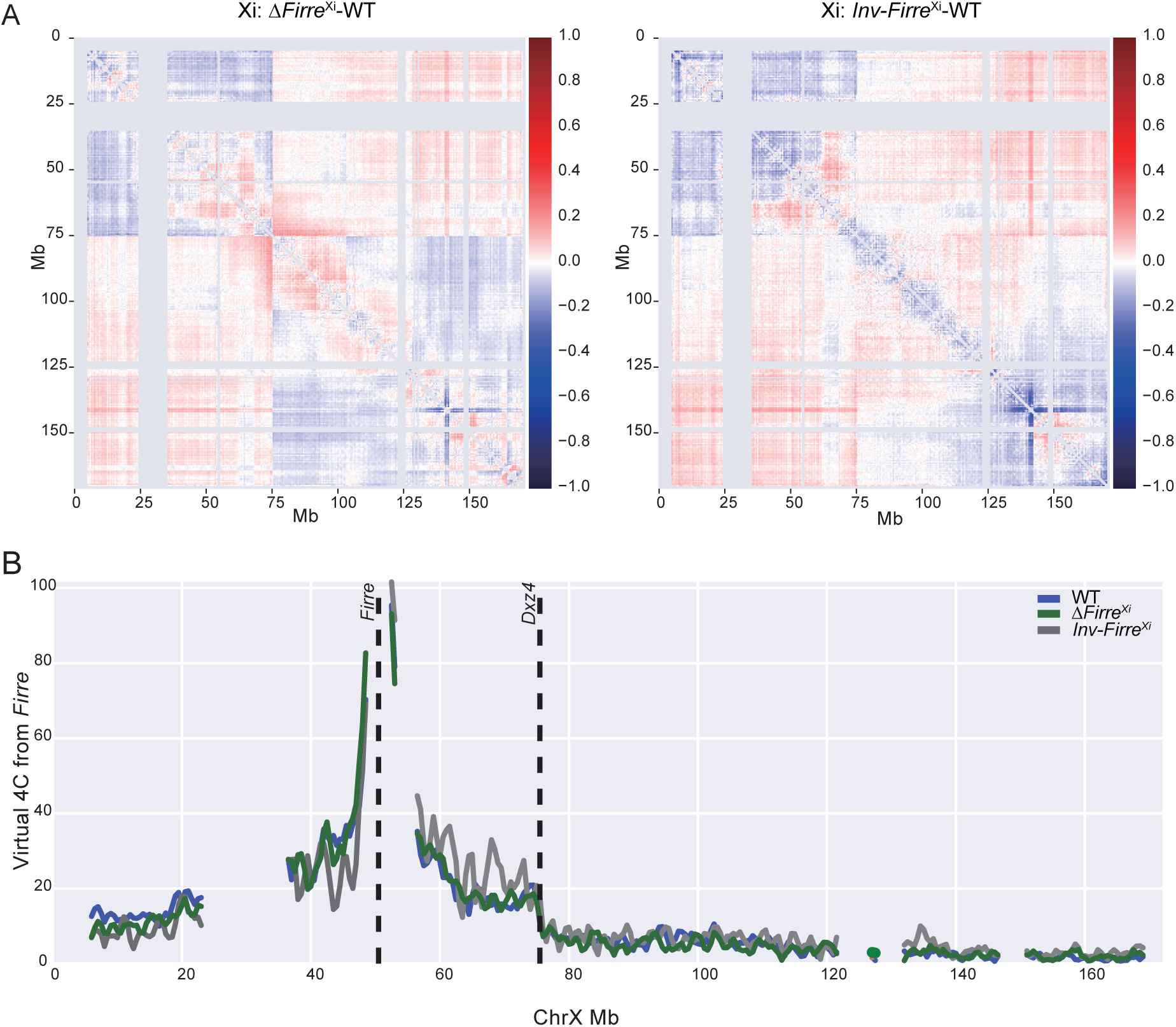
Deletion and inversion of *Firre* on the Xi causes a redistribution of contacts on the Xi consistent with *Firre/Dxz4* interactions. **A**. Pearson correlated-transformed differential contact maps of the Xi at 500kb resolution highlight differences between Δ*Firre*^Xi^ and WT, and between Inv*Firre*^Xi^ and WT. Loss and gain of contacts in the mutants versus WT appear blue and red, respectively. The color scale shows differential Pearson correlation values. **B**. Virtual 4C plots derived from Hi-C data at 500kb resolution using *Firre* as the viewpoint on the Xi in WT (blue), Δ*Firre*^Xi^ (green) and Inv*Firre*^Xi^ (grey). The positions of *Firre* and *Dxz4* is indicated. See also Figure 6C, D.

We conclude that loss of *Firre* RNA causes few changes in chromatin accessibility, but analysis of a double mutant shows cooperation between *Firre* and *Dxz4* in maintenance of condensed chromatin on the Xi. Alterations in contact distribution on the Xi upon loss of *Firre* RNA suggest that the lncRNA exerts trans-effects on the 3D structure of the Xi, potentially due to loss of H3K27me3 and of CTCF. Deletion or inversion of the silent *Firre* locus on the Xi exerts a cis-effect on the Xi structure.

## Discussion

Studies of lncRNAs support the notion that these molecules can either spread in cis from their genomic locus or localize to cellular compartments away from their own locus of transcription to perform essential functions in regulating gene expression ^38–41^. Here, we report that the lncRNA *Firre* is transcribed from the Xa and then acts in trans to maintain epigenetic features and nuclear location of the Xi. Importantly, ectopic expression of *Firre* RNA from a cDNA transgene partially rescues the loss of H3K27me3 or CTCF on the Xi and the association of the Xi to the nucleolus, supporting a trans-acting role. Loss of *Firre* RNA does not cause reactivation of the Xi, but results in dysregulation of genes implicated in cell division and development, which is also rescued by a transgene.

Interestingly, other lncRNAs play important roles in nuclear structure as they fold into higher-order structures and act in cooperation with proteins including chromatin-modifying complexes ^42^. *Xist* provides the quintessential example of a lncRNA that acts in cis and spreads along an entire chromosome to recruit a series of proteins that implement X chromosome silencing ^3,20,42^. In contrast, *Firre* acts in trans to recruit SUZ12 and possibly other components of PRC2 to maintain H3K27me3 on the Xi and potentially on other regions of the genome. Examples of trans-effects have been reported for the lncRNAs *Fendrr* essential for proper heart development and *Pint* important in cell proliferation, both of which recruit the PRC2 complex for H3K27 tri-methylation of loci located on other chromosomes ^43,44^. Another lncRNA, *Meg3*, recruits two PRC2 components, JARID2 and EZH2, to facilitate H3K27me3 deposition and repression of specific genes in trans ^45,46^. Interestingly, *Meg3* has an additional cis-activating role, by sequestration of PRC2 to prevent DNA methylation-induced repression of genes within the *Meg3-Mirg* imprinting cluster ^47,48^.

Loss of *Firre* RNA in Patski cells (derived from kidney from an 18.5dpc embryo) and MEFs (derived from a 13.5dpc embryo) causes a significant loss of H3K27me3 and of the PRC2 subunit SUZ12 enrichment on the Xi as well as a loss of nucleolar and lamina association of the Xi. We postulate that the nucleolus/lamina association of the Xi may be coupled with H3K27me3 enrichment and maintenance of heterochromatin. Supporting this notion is the previously reported finding of Xi decondensation as evidenced by sparser H3K27me3 staining when the Xi is away from the nucleolus or nuclear periphery ^49^. The nucleolus has emerged as a platform for the organization of chromatin enriched in repressive histone modifications ^17–19,50^. For example, in fibroblasts and cancer cells loss of the nucleolar protein NPM1 (nucleophosmin) results in deformed nucleoli and in redistribution of H3K27me3 ^51^. A recent study shows that deleting *Xist* also results in losses of both H3K27me3 and nucleolar association of the Xi ^52^. However, we did not observe disruption of *Xist* RNA expression nor coating on the Xi in Δ*Firre*^Xa^ cells. We thereby propose that *Firre* may have a role in maintenance of H3K27me3 and nucleolar association independent of *Xist*. This role may be in part mediated by CTCF, an important factor that facilitates interactions between genomic regions and the nucleolus ^37^. Indeed, we found a significant loss of CTCF binding on the entire Xi when *Firre* RNA is absent, which could explain abnormal anchoring of the Xi to the nucleolar periphery. Other factors may be implicated, since we found that loss of *Firre* RNA also causes downregulation of *Ezh1*, a protein implicated in H3K27 trimethylation, and of the linker histone H1 important in chromatin condensation ^53,54^.

Importantly, the loss of H3K27me3 and of CTCF and the abnormal location of the Xi could all be rescued at least in part by a mouse *Firre* cDNA transgene and to a lesser extent by a human *FIRRE* cDNA transgene. These findings imply that sequences that mediate maintenance of H3K27me3 and location of the Xi are present in the exons of the gene. Previous studies have shown that repeats (RRD) contained in *Firre/FIRRE* exons display 65% identity between the species, and that these repeats bind the nuclear matrix protein hnRNPU ^22,32^. This nuclear organizer protein is known to associate with *Xist* and the Xi, as well as to multiple other regions of the genome ^21,22,24,55–58^.

Our results are consistent with dosage effects of *Firre*, since addition of an ectopic transgene increased H3K27me3 to high levels on the Xi in all cell types tested. Overexpression of *Firre* was found to impact the frequency of natural killer cells and disrupt hematopoietic stem cell transplantation ^31,33^. Amplification of *FIRRE,* along with two neighboring genes, *IGSF1* and *OR13H1*, has been implicated in periventricular nodular heterotopia that causes epilepsy in humans, suggesting that the dosage of *FIRRE* RNA is critical to normal cellular physiology ^59^. Furthermore, high *FIRRE* expression in several tumor types, especially kidney clear cell carcinoma, brain lower grade glioma and liver hepatocellular carcinoma is associated with decreased survival rates ^60^.

Surprisingly, we did not observe a loss of H3K27me3 enrichment on the Xi in tissue sections from brain, kidney and liver from *Firre* KO mice. Similarly, Froberg and colleagues reported that mouse ES cells with a *Firre* deletion on the Xa, the Xi or on both alleles show no loss of H3K27me3 on the Xi after differentiation ^35^. Thus, *Firre* may play a role in maintenance but not in initiation of H3K27me3 enrichment on the Xi. Alternatively, these conflicting results may be explained by cell type/tissue differences, and/or by compensatory mechanisms in early development.

We found gene dysregulation in cells without *Firre* RNA, which could be rescued by a mouse *Firre* cDNA transgene. Similar to our observations unspecific expression changes in genes implicated in cell cycle and in development have been reported for other ablated lncRNAs, which may reflect changes in cell growth (Goff et al., 2015). A new study reports that *Firre* KO mice show organ-dependent autosomal gene dysregulation, with more differentially expressed genes in spleen than in other tissues ^31^. Concordantly, *Firre* KO mice have abnormalities in B- and T-cell physiology, that can be rescued by ectopic expression of *Firre* from autosomes defining a trans-acting role in vivo ^33^. In human cells, induced loss of *FIRRE* RNA causes dysregulation of inflammatory gene expression ^24^. The changes in gene expression we observed in kidney-derived Patski cells mainly affect cell cycle and development, which differ from those reported in *Firre* KO mouse spleen, suggesting tissue-specific differences and/or compensatory mechanisms during early development. Such compensatory effects in KO organisms can mask phenotypes ^61^. The minor dysregulation of X-linked genes and lack of changes in Xi structure and chromatin accessibility observed in Δ*Firre*^Xa^ cells despite H3K27me3 loss on the Xi are consistent with multiple silencing layers of XCI control, and agree with a new report of normal imprinted XCI in placenta of *Firre* KO mice ^31^. It should also be noted that H3K27me3 and gene expression changes do not necessarily correlate ^62,63^.

We found that inversion of *Firre* on the Xi results in an orientation-dependent contact redistribution, which supports an essential role for CTCF motif orientation in the formation of higher-order chromatin structure ^64^. We reported a similar but stronger effect at *Dxz4* after its inversion due to reorientation of unidirectional CTCF binding motifs ^29^. The *Firre/FIRRE* loci bind CTCF preferentially on the Xi via canonical motifs located within their introns ^21,22,65^. In contrast, the multiple CTCF motifs at the *Firre* locus have divergent orientations, with four motifs pointing to the telomere and eight to the centromere ^22^. Here, we found that inter-superdomain interactions increase in cells with a deletion of the *Firre* locus on the Xi, suggesting a role for *Firre* in helping to insulate the two superdomains perhaps via a superloop with *Dxz4* ^27^. Gain of chromatin accessibility in a double mutant line Δ*Firre*^Xi^/ΔDxz4^Xi^ support a synergistic role for the two lncRNA loci in the compaction of the Xi.

## Methods

### Cell lines and tissues

The Patski cell line, originally derived from embryonic kidney (18.5 dpc) from a cross between a female C57BL/6J (BL6) with an *Hprt^BM3^* mutation and a male *Mus spretus* (*spretus*), was previously selected in HAT (hypoxanthine-aminopterin-thymidine) medium so that the Xi is always from BL6 ^66,67^. Primary fibroblast cultures were also derived from a 13.5 dpc female F1 embryo with skewed X inactivation toward the *spretus* X chromosome due to an *Xist* mutation of the BL6 X chromosome ^68^. Cells were cultured as described and the presence of normal X chromosomes verified by karyotyping ^29^.

Tissues (liver, kidney, brain) were collected from a *Firre* KO mouse model ^33^ Tissues were collected from heterozygous and homozygous mutants and control female mice verified by genotyping. Ectopic expression of *Firre* induced by doxycycline (DOX) injection in mice was done as described ^33^ Mouse embryonic fibroblasts (MEFs) were derived from 13.5 dpc embryos of mutant (heterozygous and homozygous) and control female mice.

For ectopic *Firre* expression assays in Patski cells and in MEFs, a mouse *Firre* cDNA plasmid (mtransgene; Dharmacon BC055934) or a human *FIRRE* cDNA plasmid (htransgene; Dharmacon BC03858) were each transfected together with the selectable marker pPGK-Puro plasmid (gift from R. Jaenisch; Addgene 11349) into Δ*Firre*^Xa^ cells using Lipofectamine 3000 (Invitrogen). After transfection, Δ*Firre*^Xa+mtransgene^ or Δ*Firre*^Xa+htransgene^ were selected in Eagle’s medium with 2μg/ml puromycin for 72h, followed by recovery in Eagle’s medium with 1μg/ml puromycin for 10 days.

### Allele-specific CRISPR/Cas9 editing and RNAi knockdown

For allele-specific CRISPR/Cas9 editing of the endogenous *Firre* locus, three highly specific sgRNAs with BL6 or *spretus* SNPs at the PAM site and with low off-target scores were chosen and aligned back to the reference genome using BLAT (UCSC) to verify specificity (Supplementary Table S1). The sgRNAs cloned into px330 plasmids (Addgene) were transfected into wild-type (WT) Patski cells using Ultracruz reagents (Santa Cruz). A Patski line with a deletion of *Dxz4* was also transfected to generate a double mutant Δ*Firre*^Xi^/ΔDxz4^Xi^ ^29^. Single-cell derived colonies were selected and deletions or inversions of the targeted *Firre* locus verified using PCR and Sanger sequencing to confirm allele-specific editing (Supplementary Table S2). *Firre* RNAi knockdown was performed as described ^21^. Cells were harvested after double siRNA treatment and qRT-PCR performed to verify knockdown efficiency.

### Immunofluorescence, RNA-FISH and DNA-FISH

Immunofluorescence was done on cells grown on chamber slides, fixed with paraformaldehyde, permeabilized, and blocked as described previously ^21^. Mouse liver, kidney and brain were embedded in a cassette and sectioned by the University of Washington histopathology service center. Tissue sections (5μm) were permeabilized using 0.5% Triton X-100 for 10min and fixed in 4% paraformaldehyde for 10min. After incubation with a primary antibody specific for H3K27me3 (Upstate/Millipore, #07-449), H2AK119ubi (Cell Signaling #8240S), macroH2A (Abcam, #ab37264), or nucleophosmin (Abcam, #ab10530) overnight at 4°C in a humidified chamber cells/tissue sections were washed in 1x PBS buffer and incubated with a secondary antibody conjugated to Texas Red (anti-rabbit, Vector # TI-1000) or fluorescein (anti-mouse, Vector, # FI-2100). RNA-FISH was done using a labeled 10kb *Xist* cDNA plasmid (pXho, which contains most of *Xist* exon 1) as described ^21^. DNA-FISH was done using labeled BAC probes containing *Firre* (RP24-322N20) or *Dxz4* (RP23-299L1) ^29^.

Slides were examined by fluorescence microscopy to score the number of nuclei with enrichment in each histone modification on the Xi, using *Xist* RNA-FISH for Xi identification. A minimum of 200 nuclei were scored per cell type by at least two observers. Measurements of overall H3K27me3 staining intensity outside the Xi cluster were done in a minimum of 100 nuclei using ImageJ ^36^. To minimize the experimental variance, a mixture of 80% Δ*Firre*^Xa^ and 20% WT cells were grown in the same chamber prior to immunostaining with H3K27me3 and a control histone panH4 (Abcam ab10158) together with DNA counterstaining with Hoechst 33342. Selected nuclear areas away from the Xi were used for measurement of H3K27me3 median intensity, with same-area normalization to either Hoechst 33342 or panH4 staining. Comparisons between WT and Δ*Firre*^Xa^ were done by calculating either the H3K27me3 staining intensity versus Hoechst 33342 or panH4 for WT and Δ*Firre*^Xa^ cells separately, or the relative H3K27me3 staining intensity in Δ*Firre*^Xa^ cells versus WT cells present in the same microscope field. The location of the Xi with respect to the nuclear periphery and the edge of the nucleolus (labeled with nucleophosmin) was recorded for at least 200 cells.

### Allelic in situ DNase Hi-C, ATAC-seq, RNA-seq, ChIP-seq and CUT&RUN

In situ DNase Hi-C was done on intact nuclei from Δ*Firre*^Xa^, Δ*Firre*^Xi^, Inv*Firre*^Xi^, and WT cells as described ^29,69^. Hi-C libraries were sequenced using 150bp paired-end reads. ATAC-seq on Δ*Firre*^Xa^, Δ*Firre*^Xi^, Inv*Firre*^Xi^, Δ*Firre*^Xi^/ΔDxz4^Xi^ and WT cells, and RNA-seq on Δ*Firre*^Xa^, Δ*Firre*^Xa+mtransgene^ and WT cells were done as previously described ^29^. ChIP-seq was done on Δ*Firre*^Xa^ and WT cells using an antibody for H3K27me3 and an established protocol (Yang et al., 2015). CUT&RUN was done on Δ*Firre*^Xa^, Δ*Firre*^Xa+mtransgene^ and WT cells using antibodies for CTCF, and on Δ*Firre*^Xa^ and WT for SUZ12 (Abcam ab12073) using a published protocol ^70^. ATAC-seq, RNA-seq, ChIP-seq and CUT&RUN libraries were sequenced as 75bp pair-end reads. All sequencing datasets were analyzed to assign reads to the *spretus* or BL6 genomes using a previously developed allele-specific data analysis pipeline ^29^. Micro-RNA-seq (less than 200nt) was done in Δ*Firre*^Xa^ and WT Patski cells by BGI Genomics (https://www.bgi.com/us/). GO analysis was done using http://geneontology.org/.

## Acknowledgements

This work was supported by grants GM046883 (CMD), DK107979 (WSN and JS) from the National Institutes of Health.

## Data availability

ChIP-seq, RNA-seq, ATAC-seq and CUT&RUN datasets will be submitted to GEO.

## Supplementary material

**Supplementary Table S1**. List of sgRNAs for CRISPR/Cas9 editing

**Supplementary Table S2**. PCR primers

**Supplementary Table S3:** X-linked genes dysregulated in Δ*Firre*^Xa^ compared to WT

**Supplementary Table S4**. Predicted novel miRNAs in Δ*Firre*^Xa^ cells

**Supplementary Table S5.** Rescue efficiency with transgene in Δ*Firre*^Xa^

**Supplementary Table S6.** Autosomal genes dysregulated in Δ*Firre*^Xa^ compared to WT

**Supplementary Table S7.** Autosomal genes on chromosomes without aneuploidy dysregulated in Δ*Firre*^Xa^ cells compared to WT

**Supplementary Table S8**. GO terms for dysregulated genes in Δ*Firre*^Xa^ as compared to WT, and rescued in Δ*Firre*^Xa+mtransgene^

**Supplementary Table S9**. Summary of read counts for in situ DNase Hi-C, RNA-seq, ATAC-seq, H3K27me3 ChIP-seq, CTCF CUT&RUN and SUZ12 CUT&RUN read counts

## Supplementary Figure legends

**Supplementary Figure S1**. **No apparent change in H3K27me3 staining over the genome in Δ*Firre*^Xa^**. **A**. Box plots of the intensity of H3K27me3 staining relative to Hoechst 33342 or panH4 staining for WT (blue) and Δ*Firre*^Xa^ (red). No significant differences were seen (p = 0.8690 for Hoechst normalization; p = 0.9585 for panH4 normalization with two-tail t test). **B**. Fold change of the intensity of H3K27me3 staining in Δ*Firre*^Xa^ cells relative to the WT cells grown on the same slide and observed in the same microscope field. H3K27me3 staining was normalized either to Hoechst 33342 of to panH4. No significant deviations from a ratio of 1 were seen (p = 0.6551 for Hoechst normalization; p = 0.5986 for panH4 normalization with two-tail t test).

**Supplementary Figure S2**. **X-linked gene expression changes in Δ*Firre*^Xa^ versus WT are partially rescued by a mouse transgene**. **A**. Rescue efficiency of dysregulated, upregulated and downregulated genes in Δ*Firre*^Xa^ cells and WT. 1226 out of 1591 dysregulated genes, 652 out of 813 upregulated genes and 574 out of 778 downregulated genes are rescued or overcorrected (see Supplementary Table S5). **B.** Analysis of dysregulated genes located on autosomes without any aneuploidy in Δ*Firre*^Xa^ cells and WT. The Venn diagrams show the number of upregulated (orange) and downregulated (blue) genes in Δ*Firre*^Xa^ versus WT and in Δ*Firre*^Xa^ versus Δ*Firre*^Xa+mtransgene^. The overlapping gene set represents dysregulated genes in Δ*Firre*^Xa^ that are rescued by transgene expression. **C**. Scatter plots of allelic gene expression from the Xa and Xi for 382 expressed X-linked genes in Δ*Firre*^Xa^ and WT cells. On the Xa 28 and 15 genes are downregulated (green) and upregulated (red) >2-fold, respectively, in Δ*Firre*^Xa^. On the Xi 10 and 8 genes are upregulated (red) and downregulated (green), respectively, in Δ*Firre*^Xa^. Allelic TPM (transcripts per million) calculated from RNA-seq based on ratios of Xa/Xi SNP reads. **D**. Rescue efficiency of dysregulated genes on the Xa and on the Xi. 29 out of 43 Xa genes and 12 out of 18 Xi genes are rescued (see Supplementary Table S5). **E**. Metaplots show average H3K27me3 occupancy at genes dysregulated in Δ*Firre*^Xa^ versus WT. Unchanged, downregulated and upregulated genes are color-coded. Average enrichment is shown from the transcription start site (TSS) to the termination site (TES), with 3kb (not at scale) on either side. **F.** Example of H3K27me3 enrichment changes at 6 upregulated and 6 downregulated genes based on ChIP-seq read coverage.

**Supplementary Figure S3**. **No change in H3K36me3 and H3K4me3 enrichment after *Firre* deletion on the Xa**. **A**. Density histograms of the distribution of allelic proportions of H3K36me3 peaks (*spretus*/(*spretus* + BL6)) along the autosomes and the X chromosomes for WT (blue) and Δ*Firre*^Xa^ (red). No significant shift was observed (Wilcoxon test: −log10P = 2.58). **B**. Density histograms of the distribution of allelic proportions of H3K4me3 peaks (*spretus*/(*spretus* + BL6)) along the autosomes and the X chromosomes for WT (blue) and Δ*Firre*^Xa^ (red). No significant shift was observed (Wilcoxon tests: −log10P = 0.7).

**Supplementary Figure S4**. **No change in chromatin accessibility after *Firre* deletion or inversion on the Xi**. **A.** Density histograms of the distribution of allelic proportions (*spretus*/(*spretus* +BL6)) of ATAC peaks along the X chromosomes for WT (blue), Δ*Firre*^Xi^ (green) and Inv*Firre*^Xi^ (grey). No shift is observed (Wilcoxon test: −log10P = 7 for Δ*Firre*^Xi^ and −log10P = 1 for Inv*Firre*^Xi^). **B**. Percentages of ATAC peaks in WT (blue), Δ*Firre*^Xi^ (green) and Inv*Firre*^Xi^ (grey) along the autosomes and the X chromosomes classified as *spretus*-specific, BL6-specific, or at both.

**Supplementary Figure S5**. **Hi-C analyses of WT, Δ*Firre*^Xa^, Δ*Firre*^Xi^ and Inv*Firre*^Xi^ cell lines**. **A.** Hi-C contact maps are shown at 500kb resolution for the Xa and the Xi in WT and Δ*Firre*^Xa^. The color scale shows normalized contact counts. **B.** Differential contact maps of the Xa and Xi at 500kb resolution to highlight differences between Δ*Firre*^Xa^, Δ*Firre*^Xi^, Inv*Firre*^Xi^ and WT. Loss and gain of contacts in the Δ*Firre*^Xa^, Δ*Firre*^Xi^, Inv*Firre*^Xi^ versus WT appear blue and red, respectively. The color scale shows differential values. **C.** Pearson correlated-transformed contact maps (40kb resolution) for 4Mb around the *Firre* locus highlight persistence of the strong boundary between TADs on the Xi in WT, Δ*Firre*^Xa^, Δ*Firre*^Xi^ and Inv*Firre*^Xi^. See also Figure 6B.

## References

1. Furlan, G. et al. The Ftx Noncoding Locus Controls X Chromosome Inactivation Independently of Its RNA Products. Mol Cell 70, 462–472.e8 (2018).

2. Augui, S., Nora, E.P. & Heard, E. Regulation of X-chromosome inactivation by the X-inactivation centre. Nat Rev Genet 12, 429–42 (2011).

3. Engreitz, J.M. et al. The Xist lncRNA exploits three-dimensional genome architecture to spread across the X chromosome. Science 341, 1237973 (2013).

4. Simon, M.D. et al. High-resolution Xist binding maps reveal two-step spreading during X-chromosome inactivation. Nature 504, 465–469 (2013).

5. Heard, E. & Disteche, C.M. Dosage compensation in mammals: fine-tuning the expression of the X chromosome. Genes Dev 20, 1848–67 (2006).

6. Mira-Bontenbal, H. & Gribnau, J. New Xist-Interacting Proteins in X-Chromosome Inactivation. Curr Biol 26, R338–42 (2016).

7. Pinheiro, I. & Heard, E. X chromosome inactivation: new players in the initiation of gene silencing. F1000Res 6(2017).

8. Galupa, R. & Heard, E. X-Chromosome Inactivation: A Crossroads Between Chromosome Architecture and Gene Regulation. Annu Rev Genet 52, 535–566 (2018).

9. Zylicz, J.J. et al. The Implication of Early Chromatin Changes in X Chromosome Inactivation. Cell 176, 182–197.e23 (2019).

10. Bonora, G. & Disteche, C.M. Structural aspects of the inactive X chromosome. Philos Trans R Soc Lond B Biol Sci 372(2017).

11. Jégu, T., Aeby, E. & Lee, J.T. The X chromosome in space. Nat Rev Genet 18, 377–389 (2017).

12. Andrulis, E.D., Neiman, A.M., Zappulla, D.C. & Sternglanz, R. Perinuclear localization of chromatin facilitates transcriptional silencing. Nature 394, 592–5 (1998).

13. Barr, M.L. & Bertram, E.G. A Morphological Distinction between Neurones of the Male and Female, and the Behaviour of the Nucleolar Satellite during Accelerated Nucleoprotein Synthesis. Nature 163, 676–677 (1949).

14. Lyon, M.F. Sex chromatin and gene action in the mammalian X-chromosome. Am J Hum Genet 14, 135–48 (1962).

15. Rego, A., Sinclair, P.B., Tao, W., Kireev, I. & Belmont, A.S. The facultative heterochromatin of the inactive X chromosome has a distinctive condensed ultrastructure. J Cell Sci 121, 1119–27 (2008).

16. Zhang, L.F., Huynh, K.D. & Lee, J.T. Perinucleolar targeting of the inactive X during S phase: evidence for a role in the maintenance of silencing. Cell 129, 693–706 (2007).

17. Padeken, J. & Heun, P. Nucleolus and nuclear periphery: velcro for heterochromatin. Curr Opin Cell Biol 28, 54–60 (2014).

18. Belagal, P. et al. Decoding the principles underlying the frequency of association with nucleoli for RNA polymerase III-transcribed genes in budding yeast. Mol Biol Cell 27, 3164–3177 (2016).

19. Huang, S., Deerinck, T.J., Ellisman, M.H. & Spector, D.L. The dynamic organization of the perinucleolar compartment in the cell nucleus. J Cell Biol 137, 965–74 (1997).

20. Chen, C.K. et al. Xist recruits the X chromosome to the nuclear lamina to enable chromosome-wide silencing. Science 354, 468–472 (2016).

21. Yang, F. et al. The lncRNA Firre anchors the inactive X chromosome to the nucleolus by binding CTCF and maintains H3K27me3 methylation. Genome Biol 16, 52 (2015).

22. Hacisuleyman, E., Shukla, C.J., Weiner, C.L. & Rinn, J.L. Function and evolution of local repeats in the Firre locus. Nat Commun 7, 11021 (2016).

23. Barutcu, A.R., Maass, P.G., Lewandowski, J.P., Weiner, C.L. & Rinn, J.L. A TAD boundary is preserved upon deletion of the CTCF-rich Firre locus. Nat Commun 9, 1444 (2018).

24. Lu, Y. et al. The NF-κB-Responsive Long Noncoding RNA FIRRE Regulates Posttranscriptional Regulation of Inflammatory Gene Expression through Interacting with hnRNPU. J Immunol 199, 3571–3582 (2017).

25. Izuogu, O.G. et al. Analysis of human ES cell differentiation establishes that the dominant isoforms of the lncRNAs RMST and FIRRE are circular. BMC Genomics 19, 276 (2018).

26. Rao, S.S. et al. A 3D map of the human genome at kilobase resolution reveals principles of chromatin looping. Cell 159, 1665–80 (2014).

27. Darrow, E.M. et al. Deletion of DXZ4 on the human inactive X chromosome alters higher-order genome architecture. Proc Natl Acad Sci U S A 113, E4504–12 (2016).

28. Horakova, A.H., Moseley, S.C., McLaughlin, C.R., Tremblay, D.C. & Chadwick, B.P. The macrosatellite DXZ4 mediates CTCF-dependent long-range intrachromosomal interactions on the human inactive X chromosome. Hum Mol Genet 21, 4367–77 (2012).

29. Bonora, G. et al. Orientation-dependent Dxz4 contacts shape the 3D structure of the inactive X chromosome. Nat Commun 9, 1445 (2018).

30. Giorgetti, L. et al. Structural organization of the inactive X chromosome in the mouse. Nature 535, 575–9 (2016).

31. Andergassen, D. et al. *In vivo Firre* and *Dxz4* deletion elucidates roles for autosomal gene regulation. (2019).

32. Hacisuleyman, E. et al. Topological organization of multichromosomal regions by the long intergenic noncoding RNA Firre. Nat Struct Mol Biol 21, 198–206 (2014).

33. Lewandowski, J.P., et al. The Firre locus produces a trans-acting RNA molecule that functions in hematopoiesis. (2019).

34. Andergassen, D. et al. Mapping the mouse Allelome reveals tissue-specific regulation of allelic expression. Elife 6(2017).

35. Froberg, J.E., Pinter, S.F., Kriz, A.J., Jégu, T. & Lee, J.T. Megadomains and superloops form dynamically but are dispensable for X-chromosome inactivation and gene escape. Nat Commun 9, 5004 (2018).

36. Luense, S. et al. Quantification of histone H3 Lys27 trimethylation (H3K27me3) by high-throughput microscopy enables cellular large-scale screening for small-molecule EZH2 inhibitors. J Biomol Screen 20, 190–201 (2015).

37. Yusufzai, T.M., Tagami, H., Nakatani, Y. & Felsenfeld, G. CTCF tethers an insulator to subnuclear sites, suggesting shared insulator mechanisms across species. Mol Cell 13, 291–8 (2004).

38. Chen, L.L. Linking Long Noncoding RNA Localization and Function. Trends Biochem Sci 41, 761–772 (2016).

39. Fatica, A. & Bozzoni, I. Long non-coding RNAs: new players in cell differentiation and development. Nat Rev Genet 15, 7–21 (2014).

40. Vance, K.W. & Ponting, C.P. Transcriptional regulatory functions of nuclear long noncoding RNAs. Trends Genet 30, 348–55 (2014).

41. Engreitz, J.M. et al. Local regulation of gene expression by lncRNA promoters, transcription and splicing. Nature 539, 452–455 (2016).

42. Engreitz, J.M., Ollikainen, N. & Guttman, M. Long non-coding RNAs: spatial amplifiers that control nuclear structure and gene expression. Nat Rev Mol Cell Biol 17, 756–770 (2016).

43. Grote, P. et al. The tissue-specific lncRNA Fendrr is an essential regulator of heart and body wall development in the mouse. Dev Cell 24, 206–14 (2013).

44. Marín-Béjar, O. et al. Pint lincRNA connects the p53 pathway with epigenetic silencing by the Polycomb repressive complex 2. Genome Biol 14, R104 (2013).

45. Kaneko, S. et al. Interactions between JARID2 and noncoding RNAs regulate PRC2 recruitment to chromatin. Mol Cell 53, 290–300 (2014).

46. Yen, Y.P. et al. locus-derived lncRNAs perpetuate postmitotic motor neuron cell fate and subtype identity. Elife 7(2018).

47. Das, P.P. et al. PRC2 Is Required to Maintain Expression of the Maternal Gtl2-Rian-Mirg Locus by Preventing De Novo DNA Methylation in Mouse Embryonic Stem Cells. Cell Rep 12, 1456–70 (2015).

48. Zhao, J. et al. Genome-wide identification of polycomb-associated RNAs by RIP-seq. Mol Cell 40, 939–53 (2010).

49. Lyu, G. et al. Changes in the position and volume of inactive X chromosomes during the G0/G1 transition. Chromosome Res 26, 179–189 (2018).

50. Padeken, J. et al. The nucleoplasmin homolog NLP mediates centromere clustering and anchoring to the nucleolus. Mol Cell 50, 236–49 (2013).

51. Holmberg Olausson, K., Nistér, M. & Lindström, M.S. Loss of nucleolar histone chaperone NPM1 triggers rearrangement of heterochromatin and synergizes with a deficiency in DNA methyltransferase DNMT3A to drive ribosomal DNA transcription. J Biol Chem 289, 34601–19 (2014).

52. Colognori, D., Sunwoo, H., Kriz, A.J., Wang, C.Y. & Lee, J.T. Xist Deletional Analysis Reveals an Interdependency between Xist RNA and Polycomb Complexes for Spreading along the Inactive X. Mol Cell (2019).

53. Ezhkova, E. et al. EZH1 and EZH2 cogovern histone H3K27 trimethylation and are essential for hair follicle homeostasis and wound repair. Genes Dev 25, 485–98 (2011).

54. Kim, J.M. et al. Linker histone H1.2 establishes chromatin compaction and gene silencing through recognition of H3K27me3. Sci Rep 5, 16714 (2015).

55. Hasegawa, Y., Brockdorff, N., Kawano, S., Tsutui, K. & Nakagawa, S. The matrix protein hnRNP U is required for chromosomal localization of Xist RNA. Dev Cell 19, 469–76 (2010).

56. Pullirsch, D. et al. The Trithorax group protein Ash2l and Saf-A are recruited to the inactive X chromosome at the onset of stable X inactivation. Development 137, 935–43 (2010).

57. McHugh, C.A. et al. The Xist lncRNA interacts directly with SHARP to silence transcription through HDAC3. Nature 521, 232–6 (2015).

58. Fan, H. et al. The nuclear matrix protein HNRNPU maintains 3D genome architecture globally in mouse hepatocytes. Genome Res 28, 192–202 (2018).

59. Abe, Y. et al. Xq26.1-26.2 gain identified on array comparative genomic hybridization in bilateral periventricular nodular heterotopia with overlying polymicrogyria. Dev Med Child Neurol 56, 1221–1224 (2014).

60. Beltrán-Anaya, F.O., Cedro-Tanda, A., Hidalgo-Miranda, A. & Romero-Cordoba, S.L. Insights into the Regulatory Role of Non-coding RNAs in Cancer Metabolism. Front Physiol 7, 342 (2016).

61. Rossi, A. et al. Genetic compensation induced by deleterious mutations but not gene knockdowns. Nature 524, 230–3 (2015).

62. You, Y. et al. Temporal dynamics of gene expression and histone marks at the Arabidopsis shoot meristem during flowering. Nat Commun 8, 15120 (2017).

63. Vanhove, J. et al. H3K27me3 Does Not Orchestrate the Expression of Lineage-Specific Markers in hESC-Derived Hepatocytes In Vitro. Stem Cell Reports 7, 192–206 (2016).

64. de Wit, E. et al. CTCF Binding Polarity Determines Chromatin Looping. Mol Cell 60, 676–84 (2015).

65. Sheffield, N.C. et al. Patterns of regulatory activity across diverse human cell types predict tissue identity, transcription factor binding, and long-range interactions. Genome Res 23, 777–88 (2013).

66. Lingenfelter, P.A. et al. Escape from X inactivation of Smcx is preceded by silencing during mouse development. Nat Genet 18, 212–3 (1998).

67. Yang, F., Babak, T., Shendure, J. & Disteche, C.M. Global survey of escape from X inactivation by RNA-sequencing in mouse. Genome Res 20, 614–22 (2010).

68. Berletch, J.B. et al. Escape from X inactivation varies in mouse tissues. PLoS Genet 11, e1005079 (2015).

69. Deng, X. et al. Bipartite structure of the inactive mouse X chromosome. Genome Biol 16, 152 (2015).

70. Skene, P.J., Henikoff, J.G. & Henikoff, S. Targeted in situ genome-wide profiling with high efficiency for low cell numbers. Nat Protoc 13, 1006–1019 (2018).

