## Supplementary figures and images for "Trans- and cis-acting effects of the lncRNA *Firre* on epigenetic and structural features of the inactive X chromosome"

### Supplementary Figure 1

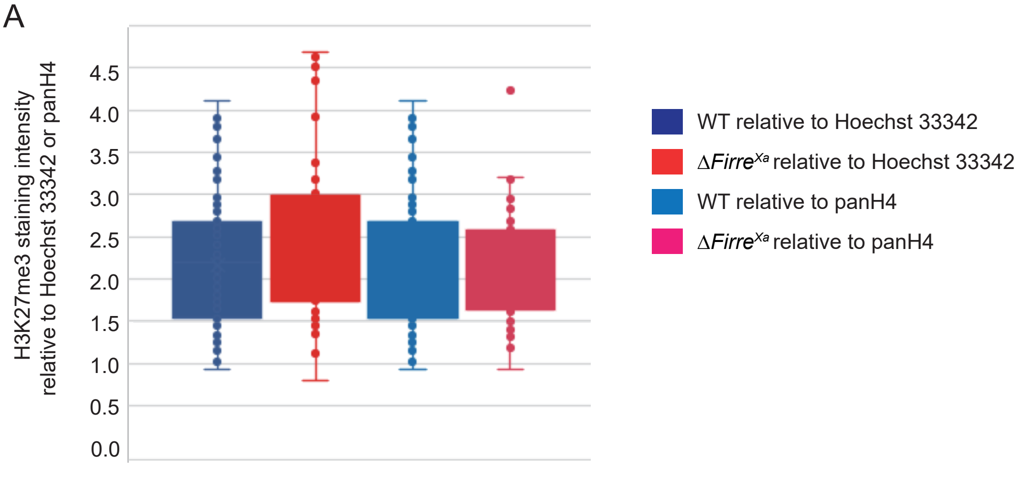

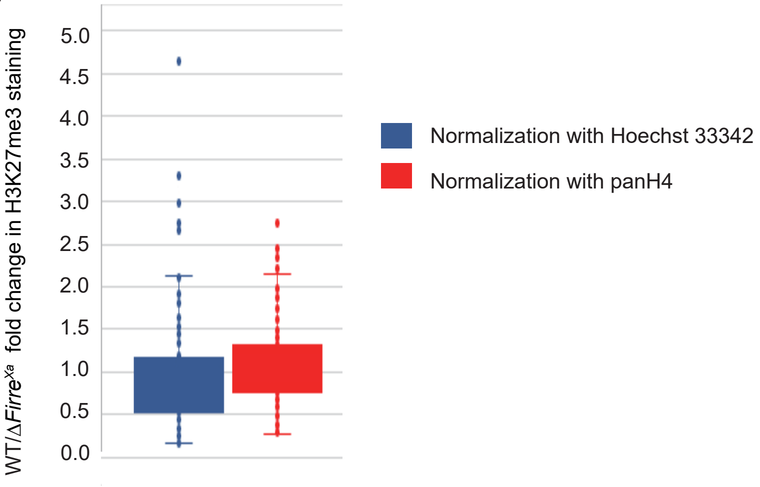

### Supplementary Figure 2

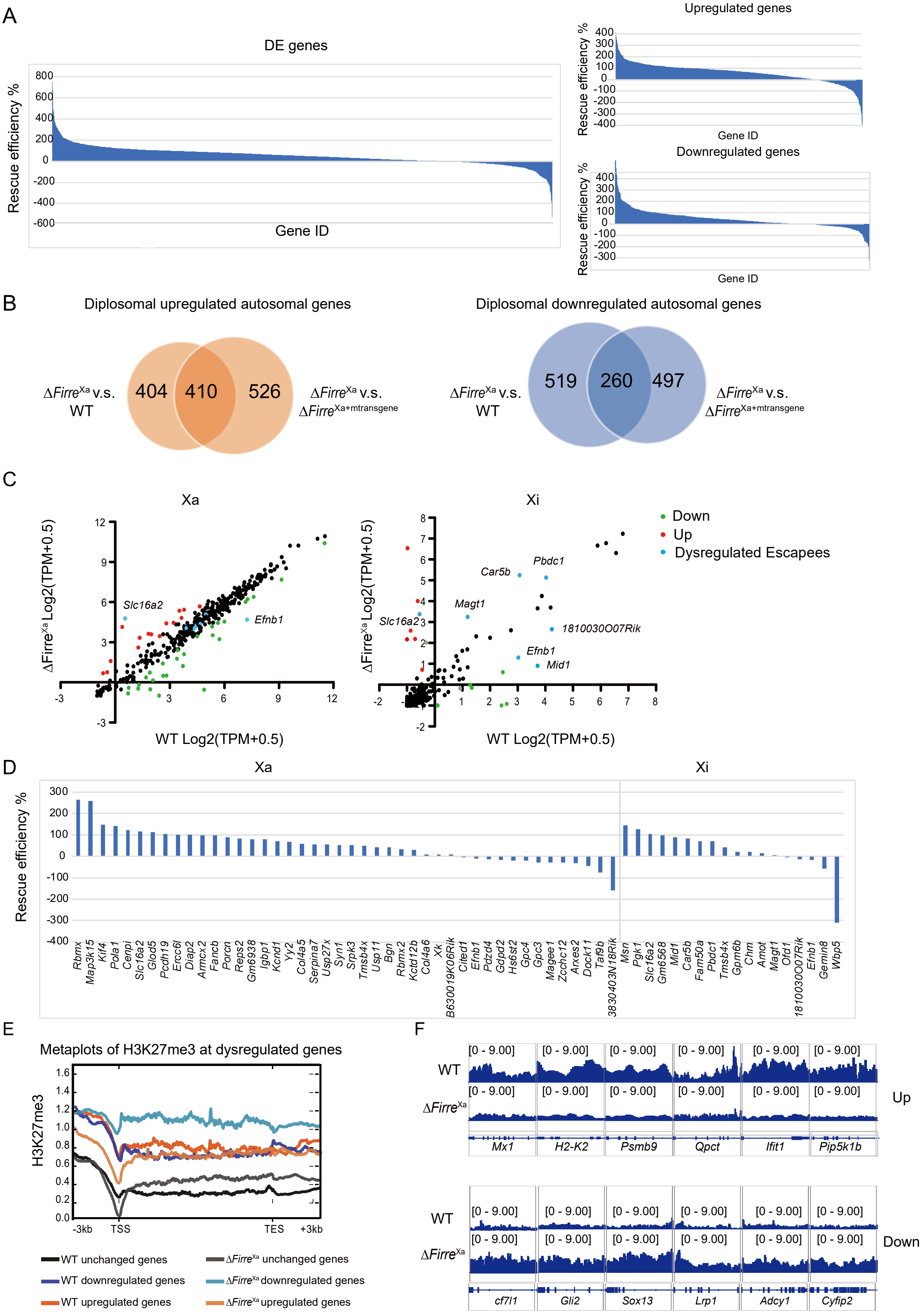

### Supplementary Figure 3

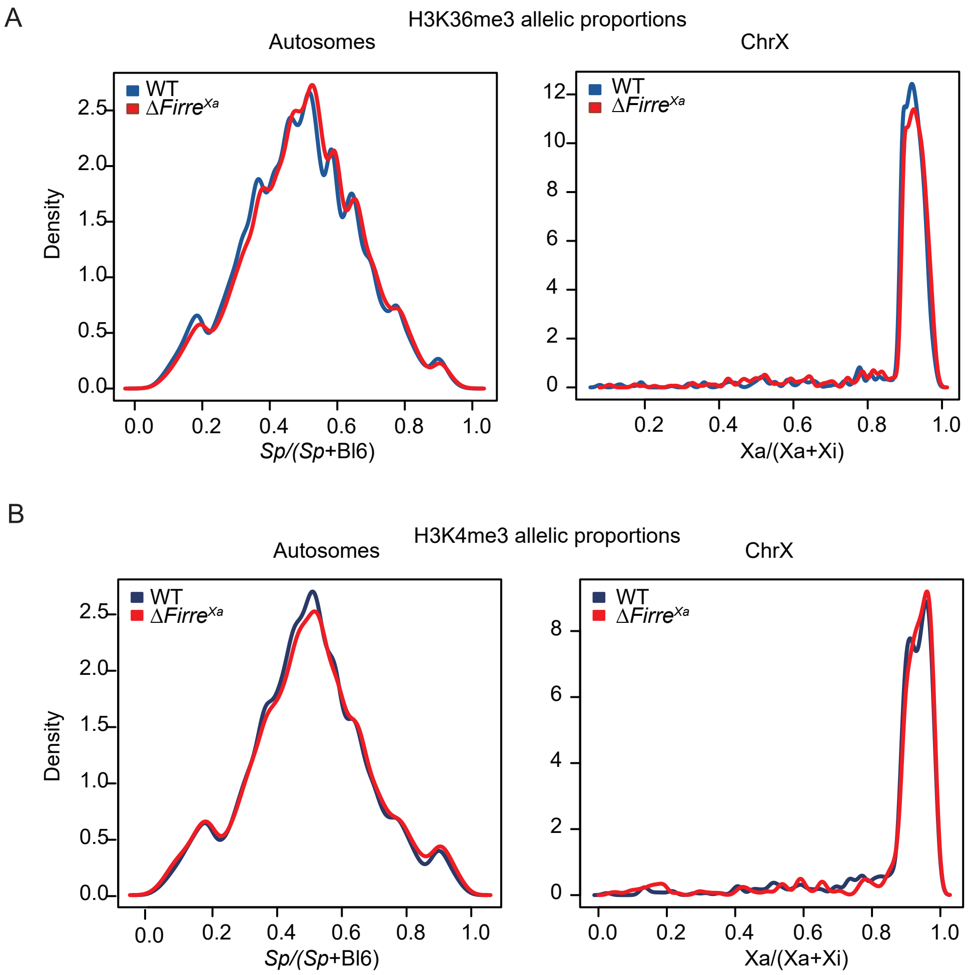

### Supplementary Figure 4

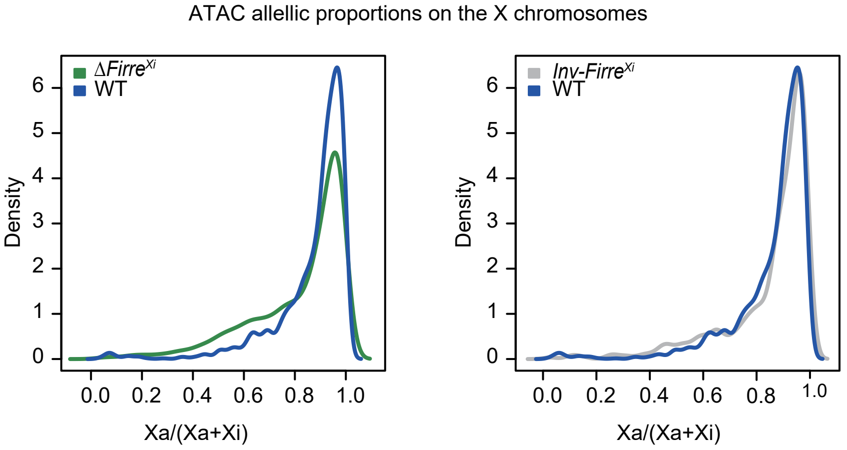

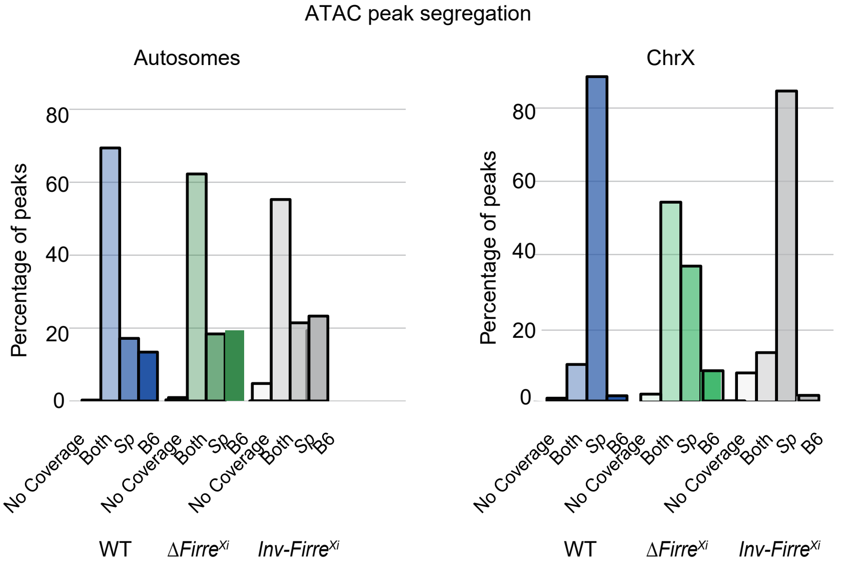

### Supplementary Figure 5

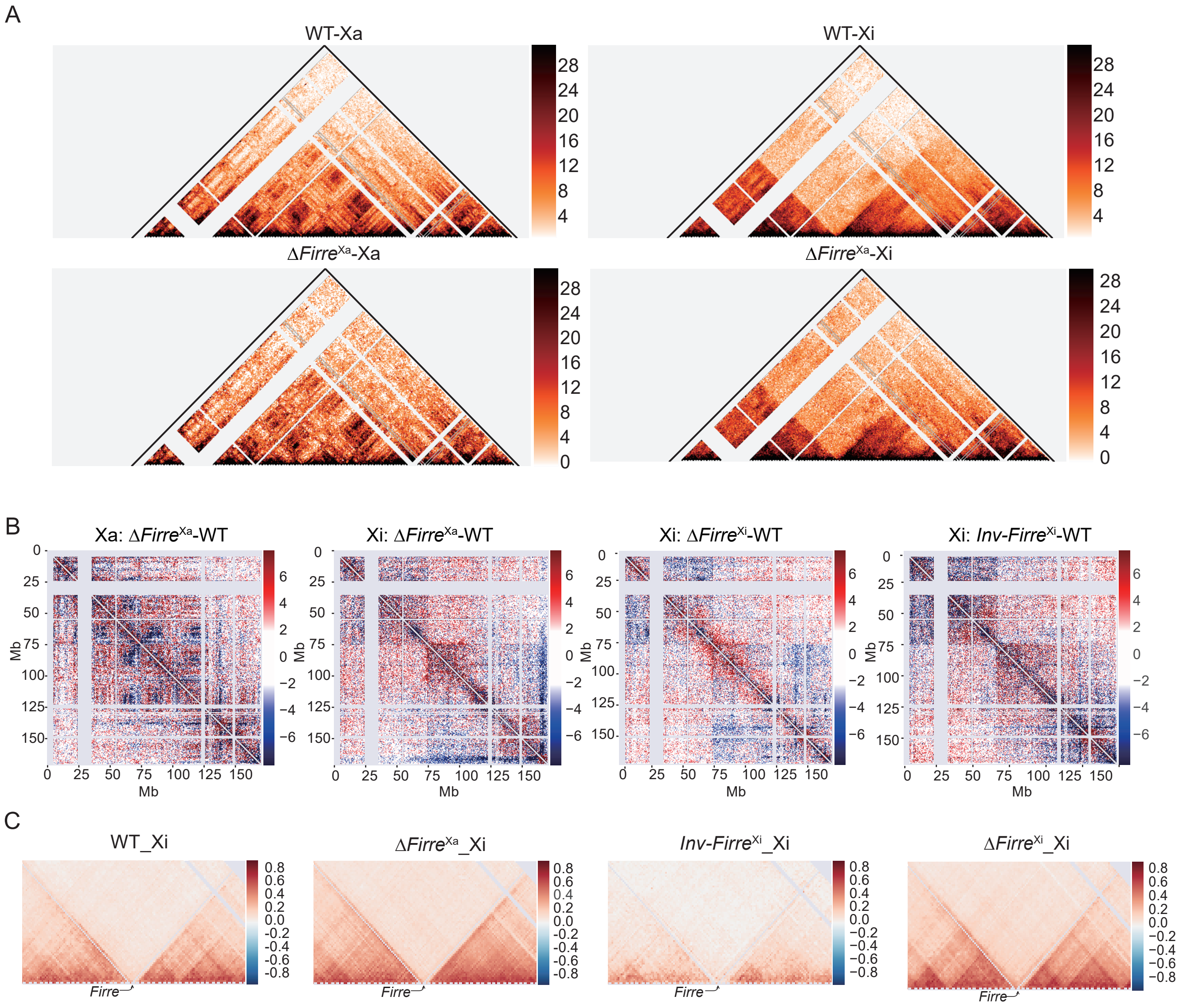
